# Chemical inhibition of a bacterial immune system

**DOI:** 10.1101/2025.02.20.638879

**Authors:** Zhiyu Zang, Olivia K. Duncan, Dziugas Sabonis, Yun Shi, Gause Miraj, Iana Fedorova, Shuai Le, Jun Deng, Yuhao Zhu, Yanyao Cai, Chengqian Zhang, Garima Arya, Breck A. Duerkop, Haihua Liang, Joseph Bondy-Denomy, Thomas Ve, Giedre Tamulaitiene, Joseph P. Gerdt

## Abstract

The rise of antibiotic resistance motivates a revived interest in phage therapy. However, bacteria possess dozens of anti-bacteriophage immune systems that confer resistance to therapeutic phages. Chemical inhibitors of these anti-phage immune systems could be employed as adjuvants to overcome resistance in phage-based therapies. Here, we report that anti-phage systems can be selectively inhibited by small molecules, thereby sensitizing phage-resistant bacteria to phages. We discovered a class of chemical inhibitors that inhibit the type II Thoeris anti-phage immune system. These inhibitors block the biosynthesis of a histidine-ADPR intracellular ‘alarm’ signal by ThsB and prevent ThsA from arresting phage replication. These inhibitors promiscuously inhibit type II Thoeris systems from diverse bacteria—including antibiotic-resistant pathogens. Chemical inhibition of the Thoeris defense improved the efficacy of a model phage therapy against a phage-resistant strain of *P. aeruginosa* in a mouse infection, suggesting a therapeutic potential. Furthermore, these inhibitors may be employed as chemical tools to dissect the importance of the Thoeris system for phage defense in natural microbial communities.

## Introduction

The spread of antibiotic-resistant bacteria is one of humanity’s greatest health threats.^1^ Bacteriophages (viruses that infect and kill bacteria) are a promising option for treating multidrug-resistant bacterial infections.^2^ However, phage resistance in pathogens is a parallel risk to antimicrobial resistance.^3^ Anti-phage immune systems are already widespread across many pathogenic bacteria, limiting the lytic efficacy of phages.^4–6^ We propose that small molecule inhibitors of anti-phage systems could be co-administered adjuvants to increase the efficacy of phage therapy against phage-resistant infections. Additionally, a selective small molecule inhibitor of an anti-phage system could “turn off” a single anti-phage defense to reveal the importance of that individual immune system for a bacterium or an entire microbial consortium within their native environments.

To identify small molecule inhibitors of anti-phage systems, we focused on the recently discovered Thoeris system. This immune system is widespread across bacteria—including human pathogens.^7^ Thoeris systems typically consist of two proteins, ThsA and ThsB. ThsB is a Toll/interleukin-1 receptor (TIR)-domain protein that produces a signal molecule after sensing phage infection^8^ (e.g., by sensing phage capsid proteins^9^). The signal molecule then binds to the effector protein ThsA, which activates ThsA to arrest phage replication and/or kill the host cell before new phage progeny are produced.^8^ Thoeris systems are classified into different types based on the domain structure of the ThsA protein.^7, 10, 11^ The type I Thoeris system encodes ThsA proteins with an N-terminal SIR2 domain and a C-terminal SLOG domain, while ThsA proteins in type II Thoeris systems comprise N-terminal transmembrane helices and a C-terminal Macro domain (**Figure 1a**).^7^ Although both types of Thoeris systems encode TIR-domain ThsB proteins, it has recently been demonstrated that the two types synthesize different signal molecules.^12^ In the type I Thoeris system, ThsB produces 1′′-3′ glycocyclic ADP ribose (gcADPR).^13, 14^ This “alarm” signal then binds to the SLOG domain of type I ThsA and activates the SIR2 domain to deplete intracellular NAD^+^, arresting phage replication (**Figure 1b**).^15^ On the other hand, ThsB in the type II Thoeris system generates a histidine-ADPR conjugate (His-ADPR).^12^ His-ADPR then binds to the Macro domain of type II ThsA, which triggers the oligomerization of ThsA at the cell membrane and stops phage replication.^12, 16^ Moreover, two other types of Thoeris systems, type III^10^ and type IV,^11^ were also recently reported.

**Figure 1.**
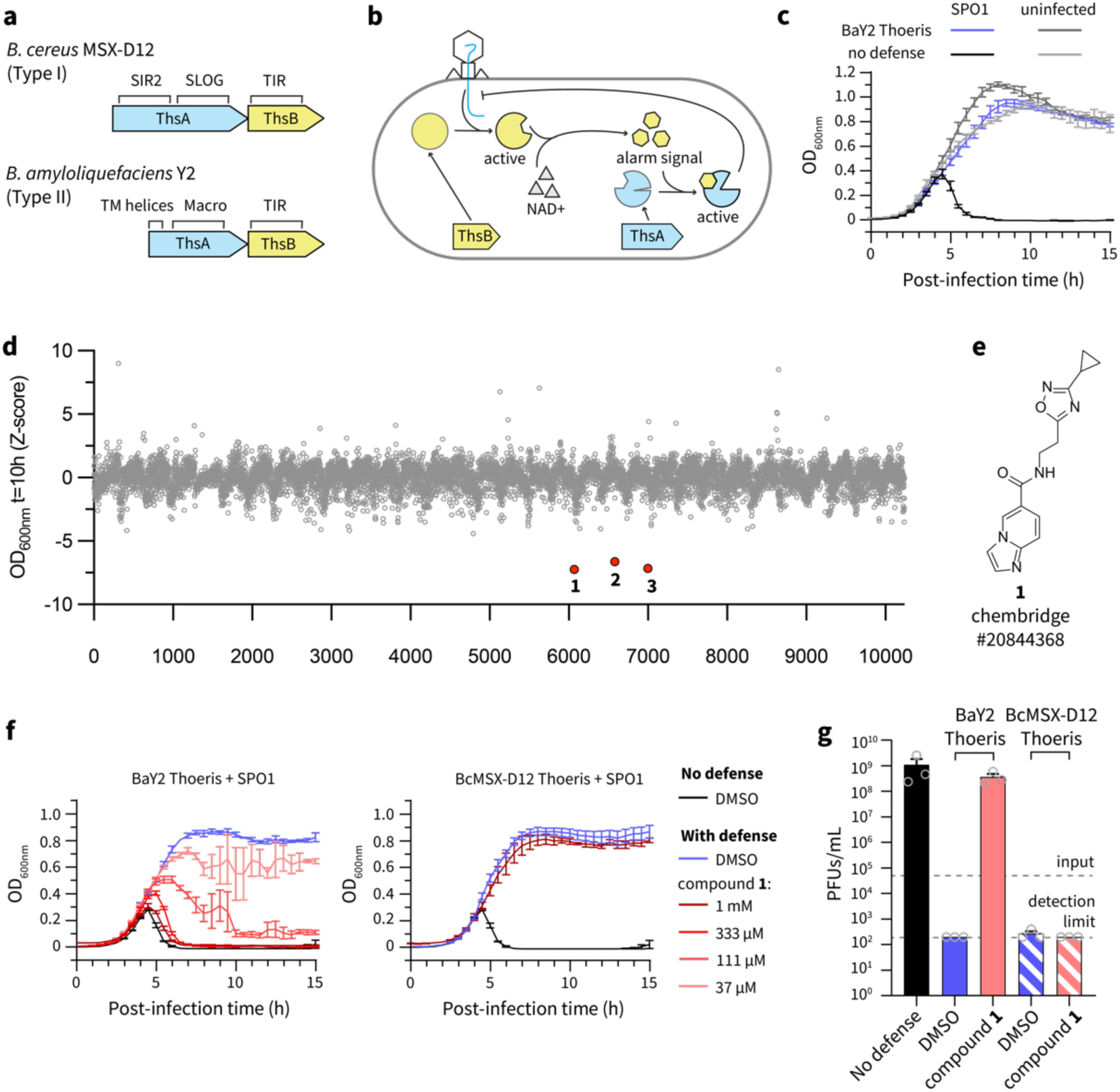
High-throughput screen identifies a specific inhibitor against type II Thoeris. (a) Gene composition (domain indicated above) of representative type I (*B. cereus* MSX-D12) and type II Thoeris (*B. amyloliquefaciens* Y2) systems. (b) Cartoon illustrating the mechanism of defense by Thoeris systems. (c) BaY2 Thoeris protection against phage infection, observed through improved phage-induced lysis in liquid culture. Where indicated, phage SPO1 was added at multiplicity of infection (MOI) of 0.01. Data are represented as the average ± SEM from three independent biological replicates. (d) Z-score plot of the screen to identify inhibitors of type II Thoeris. The Z-score is calculated based on the OD600nm difference of the compound-treated group compared to no treatment control at 10 hours post-infection. The three compounds that gave significantly lower OD600nm at 10 hours post-infection are marked red. (e) Chemical structure of compound **1**. (f) Compound **1** inhibited BaY2 Thoeris (type II) but not BcMSX-D12 Thoeris (type I) in liquid culture in a dose-dependent fashion, observed through improved phage-induced lysis in liquid culture. Data are represented as the average ± SEM from three independent biological replicates. (g) Phage reproduction of SPO1, quantified by measuring the plaque forming units (PFUs) 15 hours post-infection. Input indicates the initial PFUs in the culture. 1 mM of compound **1** was tested. Data are represented as the average ± SEM from three independent biological replicates. Each replicate is displayed with a grey circle.

Although certain phages encode proteins that inhibit Thoeris systems,^12, 13^ there are not yet any examples of small molecules that inhibit Thoeris systems. Here, we discovered a class of chemical inhibitors that specifically inhibit type II Thoeris systems. These inhibitors function by blocking the production of the His-ADPR alarm signal. We found that our inhibitors can promiscuously inhibit type II Thoeris systems in opportunistic pathogens, suggesting a therapeutic potential of these inhibitors as adjuvants to phage therapy. In vivo examination confirmed that chemical inhibition of this anti-phage defense can improve the efficacy of phage therapy in infections.

## Results

### High-throughput screen identified an inhibitor against a type II Thoeris system

To identify chemical inhibitors of Thoeris systems, the type II Thoeris operon from *Bacillus amyloliquefaciens* Y2^7^ (**Figure 1a**) was cloned into *Bacillus subtilis*. The presence of the BaY2 Thoeris system protected *B. subtilis* from SPO1 infection, as seen from the prevention of phage-induced host population lysis in liquid media (**Figure 1c**) and reduced plaque formation on solid media (**Figure S1a**). If a chemical were to inhibit the Thoeris system, we would expect the bacterial population to decrease over time due to phage-induced lysis, yielding a lower OD_600nm_. We leveraged this measurement to perform a high-throughput screen to identify chemical inhibitors of the BaY2 Thoeris system. In a screen of 10,000 synthetic compounds, 3 molecules appeared to inhibit the BaY2 Thoeris defense and enable phage-induced host population lysis (**Figure 1d, Figure S1b**). Compound **1** (**Figure 1e**) was validated to reproducibly help phage to lyse Thoeris-defended bacteria. As expected, this effect was dependent on the dose of compound **1** (**Figure 1f**) and the presence of phage **(Figure S1c**). However, neither compound **2** nor **3** reproduced a phage-dependent host population lysis. Compound **2** inhibited bacterial growth at a high concentration independently of phage infection (**Figure S1d, e**), while compound **3** failed to reproduce any detriment to the host bacteria (**Figure S1f**). Therefore, we focused on compound **1** and tested if it specifically inhibited the BaY2 Thoeris system (type II) or broadly sensitized bacteria to phages. To determine the inhibitor’s selectivity, we tested if compound **1** also inhibited a type I Thoeris system from *Bacillus cereus* MSX-D12.^7^ We cloned this Thoeris operon into *B. subtilis*, where it also afforded resistance to phage SPO1 (**Figure 1f, Figure S1a**). Notably, compound **1** did not cause a phage-induced host population lysis in the presence of the BcMSX-D12 Thoeris system (**Figure 1f**). Compound **1** also failed to accelerate the phage-induced host population lysis on a *B. subtilis* strain lacking any cloned defense (**Figure S1g**), supporting its selectivity as a type II Thoeris inhibitor. To further confirm that the observed bacterial lysis was due to compound **1** improving phage replication, we quantified phage reproduction efficiency. As expected, both the BaY2 Thoeris system and the BcMSX-D12 Thoeris system abolished phage reproduction on *B. subtilis* (**Figure 1g**), and compound **1** recovered phage reproduction only on *B. subtilis* expressing BaY2 Thoeris. As expected, compound **1** failed to increase phage replication both on *B. subtilis* expressing BcMSX-D12 Thoeris (**Figure 1g**) and *B. subtilis* cells lacking any cloned defense systems (**Figure S1h**). These results strongly suggest that compound **1** is a specific inhibitor of the type II Thoeris system.

### Structure-activity relationship study reveals inhibitors with improved potency

We then conducted a structure-activity relationship study on compound **1** (Figure 2a) to determine the necessary structural features for inhibition and to obtain the most potent inhibitor. To determine the essential components of compound **1**, we tested compound **4** (imidazo[1,2-*a*]pyridine-6-carboxamide, IP6C) and compound **5** (Figure 2b) for their defense inhibition activity relative to compound **1**. The inhibition of the Thoeris system was quantified by calculating the “Thoeris strength,” defined as the area under the lysis curve normalized to controls^17^ (**Figure S2a, Methods**). IP6C (IC_50_ = 10 µM) inhibited BaY2 Thoeris more potently than compound **1** (IC_50_ = 78 µM, Figure 2c, **Figure S2b, c**), while compound **5** (IC_50_ > 1 mM) was inactive (**Figure S2d**). As expected, IP6C also promoted the reproduction of SPO1 on *B. subtilis* cells expressing BaY2 Thoeris (Figure 2d). Therefore, IP6C (**4**) is the essential portion of compound **1** to inhibit the BaY2 Thoeris system.

**Figure 2.**
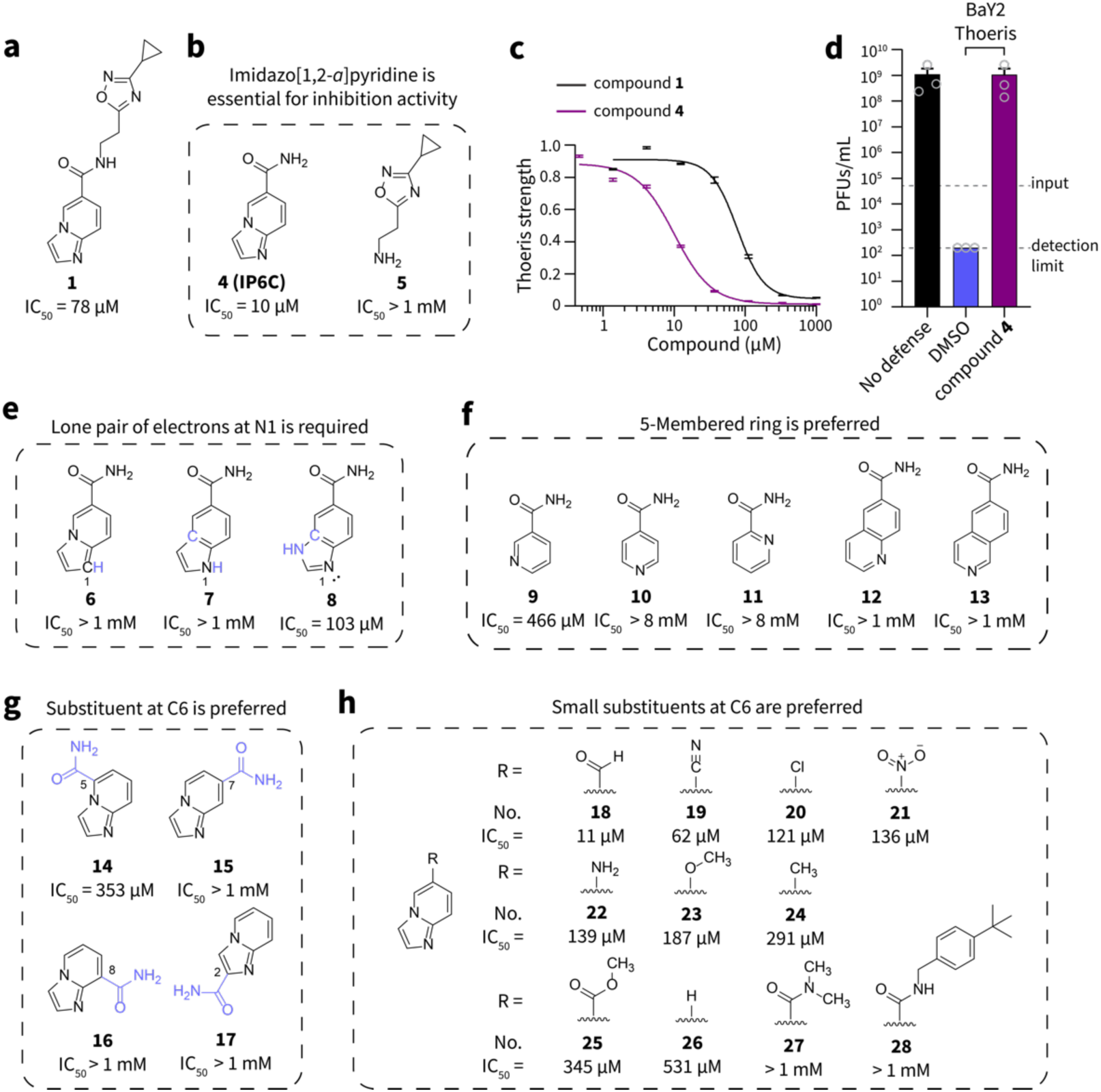
Structure-activity relationship study reveals optimal Thoeris inhibitors. (a) Chemical structure and IC50 of the initial hit (compound **1**). (b) Chemical structure and IC50 of compounds **4** – **5**. (c) Dose-response curve of compounds **1** and **4**. Data are represented as the average ± SEM from three independent biological replicates. (d) Phage reproduction of SPO1, quantified by measuring the plaque forming units (PFUs) after 15 hours post-infection. Input indicates the initial PFUs in the culture. 333 µM of compound **4** was tested. Data are represented as the average ± SEM from three independent biological replicates. Each replicate is displayed with a grey circle. (e) Chemical structures and IC50 values of compounds **6** – **8**. (f) Chemical structures and IC50 values of compounds **9** – **13**. (g) Chemical structures and IC50 values of compounds **14** – **17**. (h) Chemical structures and IC50 values of compounds **18** – **28**.

We next explored the electronics of the heterocycle to discern the necessary features for Thoeris inhibition. The imidazo[1,2-*a*]pyridine moiety contains a nitrogen atom at its 1-position with a lone pair of electrons that could either accept hydrogen bonds or act as a nucleophile. We hypothesized that the lone pair of electrons of N-1 could be important for the inhibition activity. To test this hypothesis, we examined Thoeris inhibition by compounds **6** – **8** (Figure 2e), in which the atom at the 1-position either possessed or lacked this lone pair of electrons. Indeed, when N-1 was changed to moieties lacking a basic lone pair of electrons [CH-1 (**6**) or NH-1 (**7**)], the inhibitors lost their activity (**Figure S2e, f**). However, compound **8**, which retained the lone pair of electrons at the N-1 position, remained active (**Figure S3a**). Therefore, a nitrogen at the 1-position with basic/nucleophilic electrons is essential for Thoeris inhibition.

We next asked if the imidazo[1,2-*a*]pyridine skeleton is optimal or if other sized heterocycles would be better. We tested compounds **9** – **13** (Figure 2f), which also contained a nitrogen atom with a lone pair of electrons but lacked the five-membered ring or replaced the five-membered ring with a six-membered ring. However, all adjusted skeletons were worse than IP6C (**Figure S3b – f**). Although compounds **9** – **13** share a similar skeleton with a carboxamide on the pyridine ring, only compound **9** (nicotinamide) fully inhibited the Thoeris defense (**Figure S3b**), albeit with lower potency than IP6C.

The carboxamide substituent location on the imidazo[1,2-*a*]pyridine skeleton could be another dictator of inhibition activity. To determine the optimal position, we tested compounds **14** – **17** (Figure 2g), which are isomers of IP6C but have the carboxamide at different positions on the imidazo[1,2-*a*]pyridine skeleton. None of the other positions showed improved inhibition compared to IP6C (**Figure S4a – d**), which suggested that the 6-position is the best location for the carboxamide substituent.

We finally evaluated a panel of compounds (**18** – **28**, Figure 2h) with different substituents at the 6-position. We found that small substituents at the 6-position are the most potent inhibitors (**Figure S4e – f, S5, S6**)—possibly because steric repulsion between large substituents and the target protein’s binding pocket compromises the binding affinity. An exception is the unsubstituted imidazo[1,2-*a*]pyridine (**26**): although the substituent at the 6-position (hydrogen) is the smallest among all the compounds tested, it exhibited weak potency. This observation suggests that other interactions (e.g., hydrogen bonds) between the substituent at the 6-position and the target’s binding pocket are important. Therefore, the optimal structure for the inhibitor is the imidazo[1,2-*a*]pyridine skeleton with a small substituent at the C6 position.

### Type II Thoeris inhibitors block production of the His-ADPR alarm signal

Phage defense by the type II Thoeris system involves two steps, each of which may be inhibited by IP6C. First, His-ADPR is produced by ThsB as an alarm signal upon sensing the phage infection.^12^ Subsequently, the His-ADPR signal activates ThsA to arrest phage replication in the infected host. To test if the Thoeris inhibitors block His-ADPR production by ThsB, we cloned BaY2ThsB alone onto the *B. subtilis* genome (Figure 3a). When *B. subtilis* cells expressing BaY2ThsB were infected by SPO1 phages, a new peak (*m/z* = 695.1173, negative ion mode) was detected by liquid chromatography-high resolution mass spectrometry (LC-HRMS) in the cell lysate (Figure 3b), which matched the theoretical [M−H]^-^ mass of His-ADPR. Tandem mass spectrometry (MS/MS) analysis of this peak revealed a fragmentation pattern that matched His-ADPR (Figure 3c), confirming that His-ADPR is made by BaY2ThsB upon phage infection as reported previously.^12^ The accumulation of intracellular His-ADPR was maximal 60 – 80 mins after infection by SPO1 (Figure 3d**, Figure S7a, b**), before cells were fully lysed by phages (∼90 mins post-infection). We then tested if IP6C could inhibit the production of His-ADPR by ThsB after its induction with the SPO1 phage. Indeed, IP6C abolished His-ADPR production (Figure 3d**, Figure S7c**), whereas its inactive analog did not (1H-Indole-5-carboxamide (**7**), **Figure S7d**). The loss of His-ADPR production was not due to premature cell lysis or a general depletion of cellular metabolites by IP6C because the intracellular NAD^+^ level was unchanged by IP6C treatment (**Figure S7e**). Repeated measurements of His-ADPR production at 80 mins post-infection confirmed that IP6C fully inhibited His-ADPR production in cells expressing BaY2ThsB before the cells were lysed by phages (Figure 3e**, Figure S7f**). Therefore, IP6C inhibits the type II Thoeris system by blocking His-ADPR production by ThsB.

**Figure 3.**
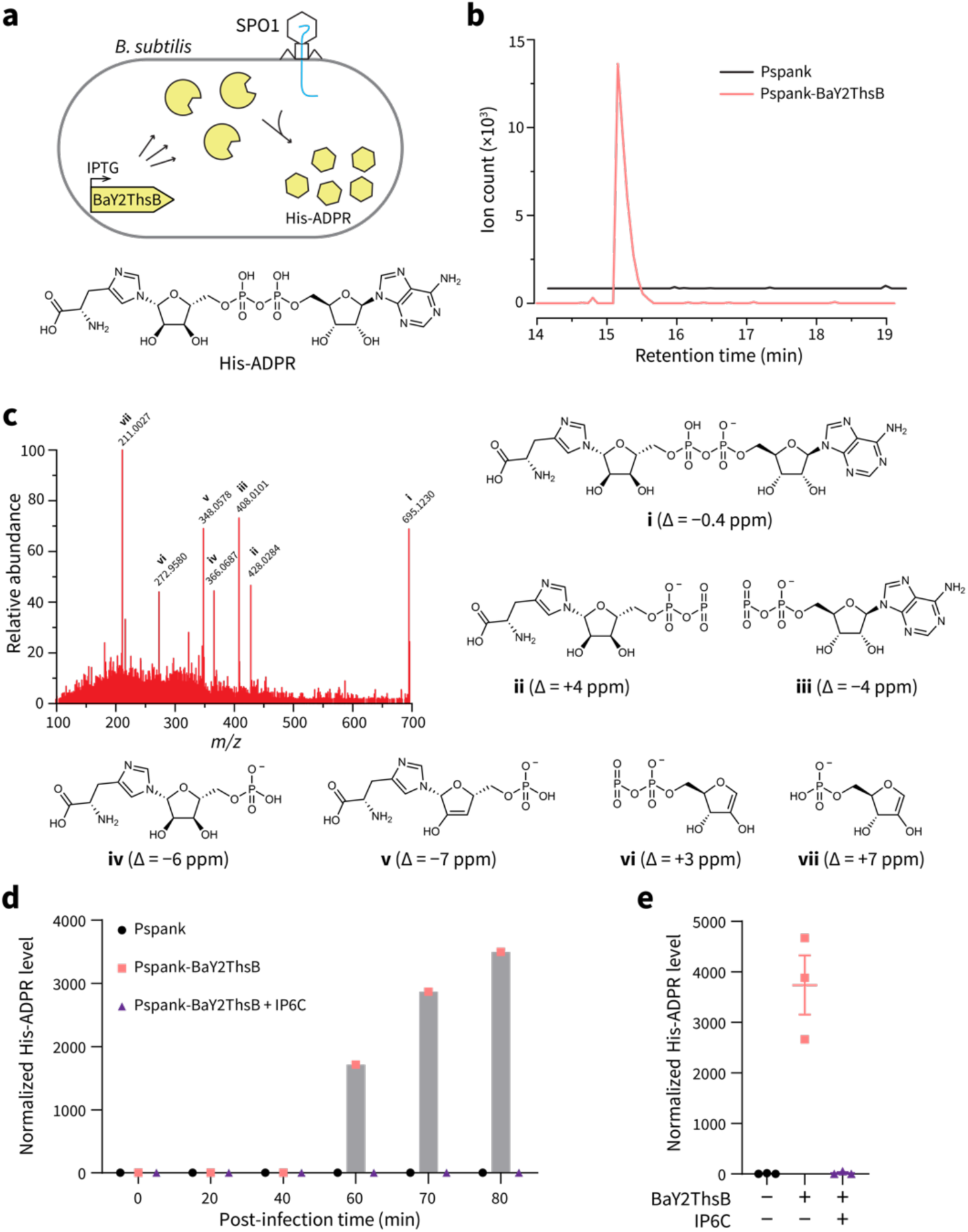
Production of the His-ADPR signal by ThsB is inhibited by IP6C. (a) Illustration of the production of His-ADPR by IPTG-induced BaY2ThsB upon phage infection. (b) Extracted ion chromatogram (EIC) of His-ADPR [*m/z* 695.1169 – 695.1212] in the lysate of cells expressing ThsB (Pspank-BaY2ThsB) or an empty promoter (Pspank) 80 mins after SPO1 infection. (c) MS/MS spectrum of His-ADPR and key fragments annotated with their associated peak number and ppm error between observed and expected *m/z* (all < 10 ppm). (d) His-ADPR level (calculated as the peak area under the EIC of His-ADPR and normalized to the standard intracellular metabolite NAD^+^) was measured at different time points following SPO1 infection. 500 µM of IP6C (**4**) was tested and DMSO was used as the negative control. (e) Biological triplicate measurement of the normalized level of His-ADPR in cell lysate 80 mins after infection with SPO1. 500 µM of IP6C was tested, and DMSO was used as the negative control. Data are represented as the average ± SEM from three independent biological replicates. Each replicate is displayed with a symbol.

### Thoeris inhibitors are competitive inhibitors of type II ThsB

The Toll/interleukin-1 receptor (TIR)-domain in the ThsB proteins is known for its NAD^+^ hydrolyzing activity.^16, 18, 19^ For example, the ThsB enzyme in the BcMSX-D12 Thoeris system (type I) converts NAD^+^ into 1′′-3′ gcADPR.^15^ It is likely that NAD^+^ and histidine are the precursors of His-ADPR synthesis by ThsB in the BaY2 Thoeris system (type II).^12^ TIR domain proteins possess a conserved glutamic acid in the catalytic pocket, which is important for their NADase activity.^18, 19^ This glutamate (Glu99) in BaY2ThsB is essential for the anti-phage activity of the Thoeris system.^7^ Therefore, we suspect that BaY2ThsB employs residue Glu99 to displace the nicotinamide from NAD^+^ and form a covalent intermediate with ADPR (Figure 4a).^19^ Then, a free histidine forms a covalent bond with ADPR, displacing the Glu99 residue to generate His-ADPR (Figure 4a**, pathway I**). Consistent with this model is our discovery that nicotinamide inhibited His-ADPR production by BaY2ThsB (**Figure S8a**), sensitizing bacteria to phages (Figure 2f**, Figure S2e**). Since nicotinamide is proposed to be the product of the initial enzymatic step, excess nicotinamide could afford product inhibition of BaY2ThsB (Figure 4a**, pathway II**).

**Figure 4.**
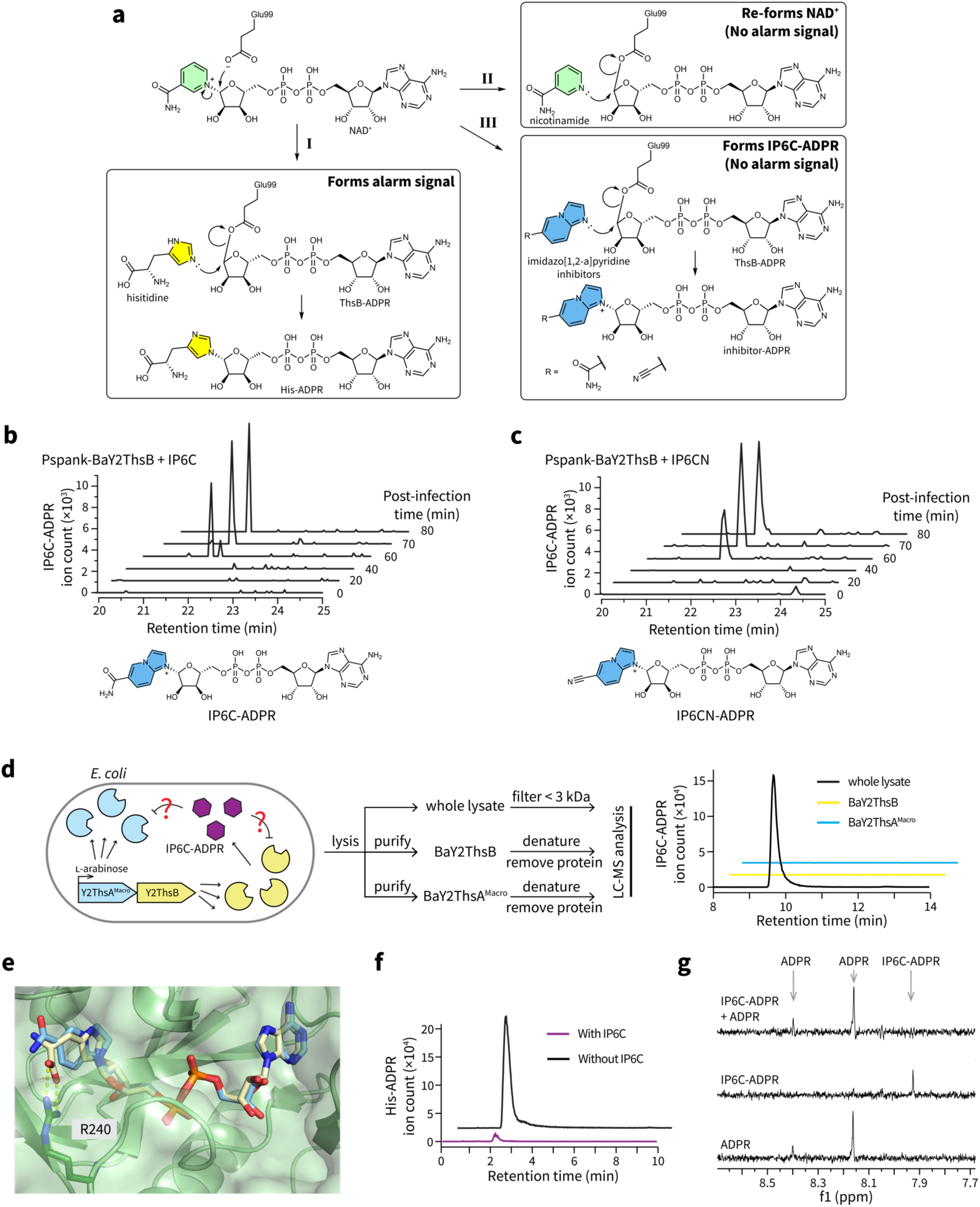
Thoeris inhibitors indirectly inhibit ThsA activation by competitively inhibiting His-ADPR production by ThsB. (a) Proposed mechanism of His-ADPR synthesis (I) and the inhibition mechanism of nicotinamide (II) and imidazo[1,2-*a*]pyridine inhibitors (III). (b) EIC of IP6C-ADPR conjugate in the lysate of SPO1-infected cells expressing BaY2ThsB. Cells were cultured with 500 µM of IP6C. (c) EIC of IP6CN-ADPR conjugate in the lysate of SPO1-infected cells expressing BaY2ThsB. Cells were cultured with 500 µM of IP6CN (**19**). (d) Scheme of IP6C-ADPR pull-down assay and EIC of IP6C-ADPR conjugate eluted from the BaY2ThsB and BaY2ThsA^Macro^ protein samples purified from cells grown with IP6C. The whole cell lysate was used as a control to verify production of IP6C-ADPR by the cells. (e) IP6C-ADPR manually docked into BaY2ThsA Macro domain complex with His-ADPR (PDB ID 8R66). The carbon atoms in the His-ADPR structure are shaded tan, and they are shaded blue in the IP6C-ADPR structure. (f) EIC of His-ADPR eluted from the BaY2ThsA^Macro^ protein sample purified from cells grown with and without IP6C (analogous to panel e, but detecting His-ADPR instead of IP6C-ADPR). (g) Expansions of STD NMR spectra showing 10 µM BaY2ThsA^Macro^ with 1 mM ADPR (bottom), 1 mM IP6C-ADPR (middle), and 1 mM ADPR + 1 mM IP6C-ADPR (top). IP6C-ADPR signal is not observed when competed with ADPR.

Because the imidazo[1,2-*a*]pyridine inhibitors may resemble histidine, we hypothesized that these inhibitors compete with histidine to bind BaY2ThsB, interfering with His-ADPR production. In fact, the nucleophilic N-1 atom in their heterocycles, which is necessary for inhibitory activity, might generate inhibitor-ADPR conjugates in a ThsB-catalyzed mechanism (Figure 4a**, pathway III**). Similar reactions exchanging heterocycle bases are catalyzed by other TIR domain enzymes.^19^ To test this hypothesis, we searched our LC-HRMS data for evidence of inhibitor-ADPR conjugates. We examined the lysates from cells that expressed BaY2ThsB and were infected by SPO1 in the presence of IP6C. Indeed, in our prior conditions when His-ADPR production was inhibited by IP6C (Figure 3d**, Figure S7c**), a new peak (*m/z* = 701.1094, negative mode) appeared in the cell lysate (Figure 4b). This peak matched the theoretical [M^+^−2H]^-^ mass of the hypothesized IP6C-ADPR conjugate. As further confirmation, we tested the analogous inhibitor compound **19** (IP6CN), which also inhibited His-ADPR production (**Figure S8b**). Similarly, a new peak (*m/z* = 683.1041, negative mode) appeared in the lysate (Figure 4c), matching the expected [M^+^−2H]^−^ mass of the IP6CN-ADPR conjugate. To verify the identity of the IP6C-ADPR generated in cells, we compared it with a purified IP6C-ADPR standard (**Figure S9a** and **Table S4**) generated via a reported enzyme-catalyzed base-exchange method^16^ in a co-injection experiment. The IP6C-ADPR made by cells expressing BaY2ThsB co-eluted with the IP6C-ADPR standard (**Figure S9b**), suggesting that they are structurally identical. Therefore, IP6C is connected to the C-1′′ position in ADPR through its N-1 atom (**Figure S9a**), as hypothesized (Figure 4a **– c**).

To validate that IP6C-ADPR was generated by BaY2ThsB alone, we assessed BaY2ThsB activity in vitro. We found that BaY2ThsB increased the rate of the formation of IP6C-ADPR from IP6C and NAD^+^ (**Figure S9c**). Admittedly, the catalysis was weak—presumably because ThsB requires activation by a phage component for robust activity. We also observed that IP6C-ADPR production was catalyzed by a BaY2ThsB homolog from *Agathobacter rectalis* ATCC 33656 (ArThsB, **Figure S9d**). A catalytically dead E99A mutant of ArThsB failed to improve IP6C-ADPR production (**Figure S9d**), further supporting our mechanistic model of IP6C-ADPR production by ThsB (Figure 4a).

Previous studies involving the human TIR domain enzyme SARM1 showed that a series of heterocyclic inhibitors were “prodrugs”, and the true SARM1 inhibitors were heterocycle-ADPR conjugates produced by SARM1.^19, 20^ We hypothesized that our imidazo[1,2-*a*]pyridine family inhibitors could also be prodrugs, and the inhibitor-ADPR conjugates produced by BaY2ThsB might be the true orthosteric inhibitors of ThsB. To test this hypothesis, we assessed if IP6C-ADPR remained bound to BaY2ThsB protein that had been purified from cells expressing BaY2ThsB in the presence of IP6C. However, although IP6C-ADPR was present in the cell lysate, it did not co-purify with BaY2ThsB (Figure 4d), suggesting that IP6C-ADPR is not a tight-binding orthosteric inhibitor of ThsB. Collectively, these results suggest that imidazo[1,2-*a*]pyridine family inhibitors are competitive inhibitors of histidine in the BaY2ThsB catalytic pocket. They cause the cell to produce inhibitor-ADPR conjugates instead of the His-ADPR alarm signal that is required to activate ThsA (Figure 4a).

### Thoeris inhibitors indirectly inhibit the activation of ThsA

Since the His-ADPR alarm signal must bind to ThsA to activate its anti-phage function,^12^ we hypothesized that IP6C and its analogs prevent ThsA activation indirectly by inhibiting His-ADPR production. However, we were also curious if the ThsB-produced inhibitor-ADPR conjugates had any direct impact on ThsA activation. For example, the inhibitor-ADPR conjugates could also serve as competitive inhibitors to prevent binding of low concentrations of His-ADPR to ThsA. Docking of IP6C-ADPR into the His-ADPR pocket of BaY2ThsA revealed that IP6C-ADPR fits well in the pocket and lacks interactions with R240 (Figure 4e), a residue that is important for ThsA activation.^12^ Therefore, IP6C-ADPR could be a competitive inhibitor of His-ADPR binding to ThsA. To test this hypothesis, we assessed the binding of IP6C-ADPR to the ThsA Macro domain. If IP6C-ADPR binds tightly to the ThsA Macro domain, we should detect IP6C-ADPR co-purified with BaY2ThsA^Macro^, as has been reported for His-ADPR binding to BaY2ThsA^Macro^.^12^ We co-expressed BaY2ThsA^Macro^ and BaY2ThsB in *E. coli* cells grown in the presence of IP6C. We then purified BaY2ThsA^Macro^ and attempted to detect IP6C-ADPR after denaturing the protein. As before,^12^ we detected His-ADPR bound to BaY2ThsA^Macro^ in the absence of IP6C. The amount of bound His-ADPR was dramatically decreased by IP6C, presumably because it prevented the production of His-ADPR by Y2 ThsB (Figure 4f). However, no IP6C-ADPR was detected in the purified BaY2ThsA^Macro^ proteins even though IP6C-ADPR was present in the cell lysate (Figure 4d). Saturation transfer difference (STD) NMR experiments also showed that IP6C-ADPR binds only very weakly to BaY2ThsA^Macro^ in vitro (Figure 4g). Notably, no binding between IP6C-ADPR and BaY2ThsA^Macro^ was detected when IP6C-ADPR was competed with an equimolar concentration of ADPR, which itself is only expected to weakly bind ThsA.^16^

Collectively, our data suggest that the inhibitor-ADPR conjugates do not bind ThsA strongly and therefore are unlikely to directly inhibit (or activate) ThsA. Instead, IP6C and its analogs indirectly inhibit the activity of ThsA by preventing the production of His-ADPR to the threshold concentration required to activate ThsA.

### Y2 Thoeris inhibitors inhibit type II Thoeris systems in opportunistic pathogens

Thoeris anti-viral systems are widespread in bacteria.^7^ Since our inhibitors generally compete with histidine binding to ThsB, we hypothesized that these inhibitors would broadly arrest BaY2-like (i.e., type II) Thoeris systems. Most importantly, we asked if our inhibitors could block type II Thoeris systems present in human pathogenic bacterial strains that are potential targets for phage therapy (e.g., multidrug-resistant strains of *Pseudomonas aeruginosa* and *Enterococcus faecalis*, Figure 5a).^21, 22^ If the inhibitors work, they could re-sensitize these phage-resistant pathogens to phage therapy. To study the efficacy of our Y2 Thoeris inhibitors on these homologous type II Thoeris systems, we cloned the Thoeris operon from the antibiotic-resistant *P. aeruginosa* clinical isolate MRSN11538^23^ and transferred it into the genome of *P. aeruginosa* PAO1, and the Thoeris operon from the antibiotic-resistant *E. faecalis* clinical isolate DS16^24^ into the genome of *E. faecalis* OG1RF. In both cases, the Thoeris systems successfully protected the bacterial hosts against phage infections (Figure 5b). Both IP6C (IC_50_ = 98 µM) and nicotinamide (IC_50_ = 2.6 mM) inhibited the type II Thoeris system in *P. aeruginosa*, re-enabling phage-induced host population lysis (Figure 5c, d**, Figure S10**). The inhibitors likewise worked in *E. faecalis* (Figure 5c, d**, Figure S10**), albeit with altered potency (IP6C IC_50_ = 8.2 mM; nicotinamide IC_50_ = 1.5 mM). We also tested if IP6C and nicotinamide could improve phage proliferation despite the presence of Thoeris systems. We found that either IP6C or nicotinamide treatment allowed both *P. aeruginosa* and *E. faecalis* phages to propagate on hosts containing type II Thoeris defense systems (Figure 5g, h). The lower potency of IP6C against *E. faecalis* could indicate weaker binding to the *E. faecalis* DS16 ThsB enzyme (which shares only 32% sequence identity with *B. amyloliquefaciens* Y2 ThsB). Other explanations are also plausible (e.g., IP6C may not permeate into *E. faecalis* well). Nonetheless, these results suggest that the Thoeris inhibitors are broad-spectrum inhibitors against multiple homologs in the type II Thoeris defense family.

**Figure 5.**
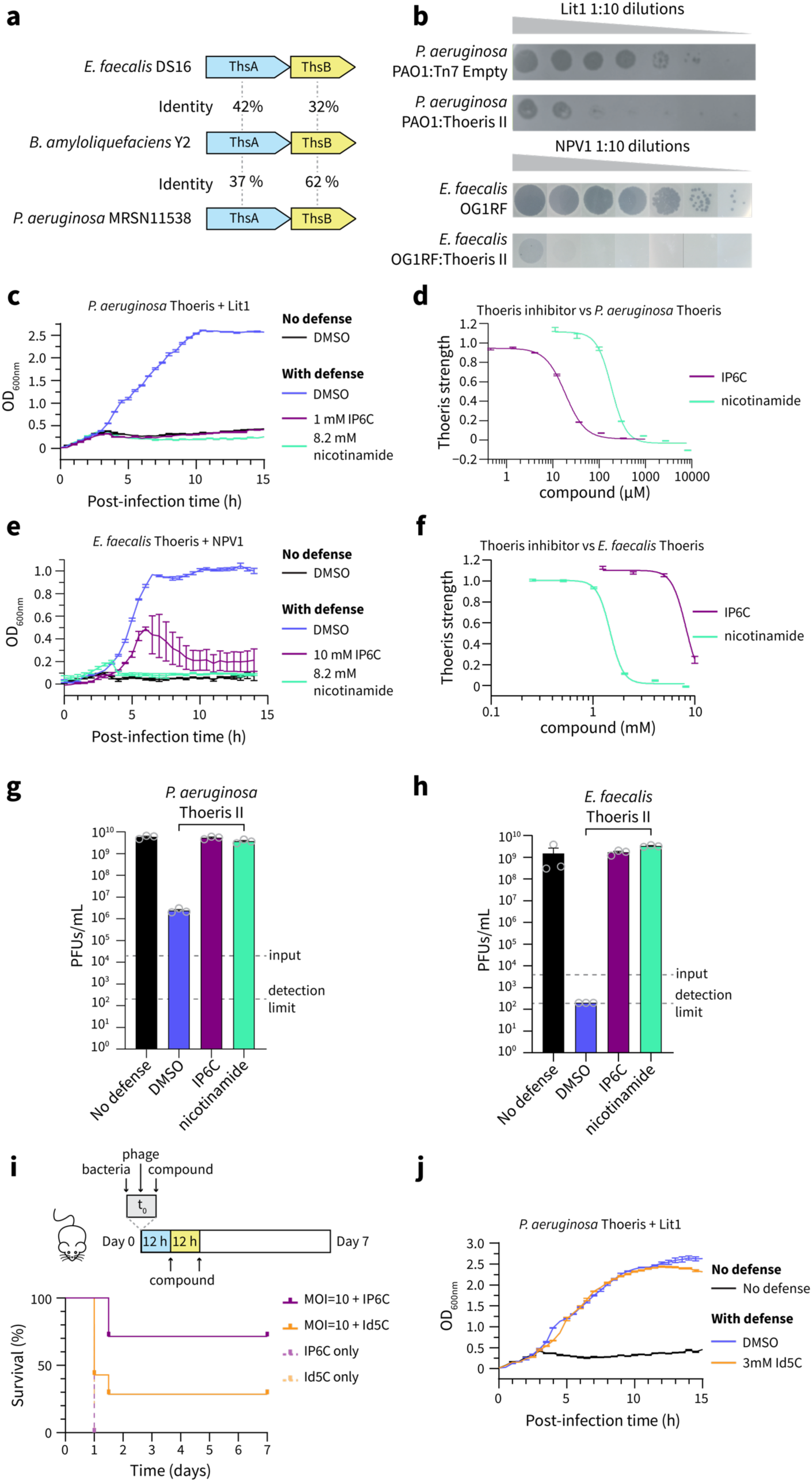
IP6C and nicotinamide inhibit type II Thoeris systems of antibiotic-resistant pathogens. (a) Percent identity between BaY2 Thoeris and two homologs present in antibiotic-resistant pathogens. (b) Type II Thoeris systems protect the pathogenic hosts from phage infection, observed through decreased plaquing on solid media by the phages Lit1 and NPV1. (c) IP6C and nicotinamide inhibit the Thoeris system in *P. aeruginosa*, observed through improved phage-induced lysis in liquid culture (MOI = 0.0001). Data are represented as the average ± SEM from three independent biological replicates. (d) Dose-response curves of Thoeris inhibition by IP6C and nicotinamide in *P. aeruginosa*. Data are represented as the average ± SEM from three independent biological replicates. (e) IP6C and nicotinamide inhibit the Thoeris system in *E. faecalis*, observed through improved phage-induced lysis in liquid culture (MOI = 0.001). Data are represented as the average ± SEM from three independent biological replicates. (f) Dose-response curves of Thoeris inhibition by IP6C and nicotinamide in *E. faecalis*. Data are represented as the average ± SEM from three independent biological replicates. (g) Phage reproduction of Lit1 on *P. aeruginosa*, quantified by measuring the PFUs 15 hours post-infection. Input indicates the initial PFUs in the culture. 1 mM of IP6C and 8.2 mM of nicotinamide were tested. Data are represented as the average ± SEM from three independent biological replicates. Each replicate is displayed with a grey circle. (h) Phage reproduction of NPV1 on *E. faecalis*, quantified by measuring the PFUs 15 hours post-infection. Input indicates the initial PFUs in the culture. 10 mM of IP6C and 8.2 mM of nicotinamide were tested. Data are represented as the average ± SEM from three independent biological replicates. Each replicate is displayed with a grey circle. (i) Survival of 7-week-old BALB/c mice (n = 7) following intraperitoneal injection with *P. aeruginosa* and three doses of compounds (each 50 mg/kg), with and without Lit1 phage at MOI=10. (j) Compound **7**, Id5C, is inactive against the type II Thoeris system in *P. aeruginosa,* observed through the lack of improved phage-induced lysis in liquid culture (even at high concentration, 3 mM [MOI = 0.0001]). Data are represented as the average ± SEM from three independent biological replicates.

### Thoeris inhibition improves the efficacy of a model phage therapy

To further evaluate the therapeutic potential of Thoeris inhibitors, we tested the ability of IP6C to improve the efficacy of a model phage therapy. We hypothesized that IP6C could improve the survival rate of mice undergoing phage therapy against phage-resistant *P. aeruginosa* containing the type II Thoeris system. In brief, mice were infected intraperitoneally with *P. aeruginosa*. The mice then received one intraperitoneal dose of Lit1 phage at a MOI of 10 and three intraperitoneal doses of IP6C inhibitor (no toxicity observed for mice, **Figure S11**) every 12 hours after administering phages (Figure 5i). We used an inactive analog of IP6C, compound **7** (Id5C, no toxicity observed for mice, **Figure S11**), as the negative control (Figure 5j). The IP6C treatment improved the survival rate of mice relative to the negative control [71% (5 out of 7) vs 29% (2 out of 7)]. The increased survival rate was dependent on phage, as all mice died within one day in the group treated with IP6C only (Figure 5i). The results of this preliminary infection model suggest that IP6C (and likely other inhibitors of anti-phage immune systems) could re-sensitize phage-resistant bacteria to phage therapies that would otherwise be ineffective.

## Discussion

We discovered a class of imidazo[1,2-*a*]pyridine derivatives that inhibits type II Thoeris systems by blocking His-ADPR production by ThsB. This finding demonstrates that anti-phage systems can be selectively inhibited by small molecules, sensitizing phage-resistant bacteria to phages. Our inhibitors arrest type II Thoeris systems in multiple bacterial species, including two multidrug-resistant opportunistic pathogens. One inhibitor improved the survival rate of mice that received phage therapy treatment to combat a phage-resistant strain of *P. aeruginosa*. Therefore, this class of inhibitors may hold future application as a therapeutic adjuvant to increase the efficacy of phage therapy against phage-resistant infections.

Apart from inhibiting type II Thoeris, this work is a blueprint to target dozens of other phage defense systems.^4^ In the coming years, selective inhibitors will likely be developed and applied against many of the most important known anti-phage immune systems. Like type II Thoeris, many anti-phage systems rely on small-molecule signaling (mostly nucleotide derivatives),^25^ such as type I Thoeris,^8, 15^ type III CRISPR,^26–28^ CBASS,^29, 30^ and PYCSAR.^31^ Among these systems, the catalytic sites of signal-synthesizing enzymes and the signal-binding sites of the effector proteins should provide deep cavities that are favorable for binding small-molecule inhibitors. On the other hand, many other anti-phage systems function through protein-protein interactions or protein-nucleic acid interactions (e.g., restriction-modification systems,^32^ CRISPR-Cas systems,^33^ Gabija,^34, 35^ Hachiman,^36^ and Zorya^37^). These types of interactions involving large interfaces (1,000–2,000 Å^2^ per side) are recalcitrant to inhibition by small molecules, but have recently proven to also be “druggable”.^38, 39^ For example, in vitro inhibitors have been developed against a CRISPR-Cas system, although they have no efficacy within bacterial cells.^40, 41^ We hypothesize that chemical inhibitors will exist for many, if not all, of the anti-phage systems. Each new inhibitor will expand the potential of phage therapy to target diverse phage-resistant infections.

Beyond the therapeutic potential of phage-defense inhibitors, they could also be useful chemical tools to dissect the importance of individual defense systems in shaping microbial communities. Many bacteria harbor multiple anti-phage systems, and the importance of each system for resistance to phages is not yet clear.^42^ To answer this question, selective inhibitors could easily ‘turn off’ individual defenses to reveal the importance of each for phage-resistance—even in genetically-intractable bacteria. Furthermore, in microbial communities, the complex benefit/cost tradeoff of harboring anti-phage immune systems^43^ promotes frequent gain and loss of anti-phage systems in individual bacteria.^44^ This constant flux of defense systems within microbial communities creates a “pan-immunity” to combat diverse phage predators and shape the composition of multi-species communities.^45^ Selective inhibitors of anti-phage systems could be easily employed to reveal the importance of individual defense systems to the “pan-immunity” of complex microbial communities. For example, an inhibitor can ‘switch off’ all type II Thoeris systems harbored by any member within a natural polymicrobial community, and the subsequent change in community composition would reveal the importance of that defense for community structure in the presence of native phages.

Finally, besides finding synthetic Thoeris inhibitors, we discovered that a natural metabolite (nicotinamide) inhibits ThsB, as well. This observation intersects with previous work showing that microbial natural products can either sensitize nearby competitors to phage lysis^46^ or provide improved resistance against phages.^47^ Similarly, our finding suggests that a microbe that secretes nicotinamide or nicotinamide-containing analogs^48, 49^ may sensitize Thoeris-containing competitors to phages. A nicotinamide-rich host environment may also preclude the effectiveness of Thoeris-based immunity. Perhaps the conditional efficacy of Thoeris (and other defenses) in certain metabolic environments is one reason why bacteria tend to maintain multiple immune systems.^42, 50^

In conclusion, we discovered a class of chemical inhibitors that can inhibit type II Thoeris anti-phage immune systems by preventing the synthesis of “alarm” signals. We demonstrated that these inhibitors work against the type II Thoeris systems encoded by multiple bacteria species, including two multidrug-resistant opportunistic pathogens. Notably, the inhibitor IP6C is also effective in vivo, where it improved phage therapy efficacy in a *P. aeruginosa*-infected mouse model. We expect that similar efforts will succeed in discovering inhibitors against dozens of other known anti-phage systems, expanding the scope of infections that can be treated with phages. We further anticipate that selective inhibitors will prove to be valuable chemical tools to study the importance of individual anti-phage systems in complex microbiomes.

## Methods

### Strains and growth conditions

The strains, bacteriophages, and plasmids used in this study are listed in **Table S1**. All chemicals used in this study are listed in **Table S2**. All primers used in this study are listed in **Table S3**. *B. subtilis* strains were routinely grown in LB broth at 37 °C or 30 °C and 220 rpm. *E. coli* and *P. aeruginosa* strains were routinely grown in LB broth at 37 °C and 220 rpm. *E. faecalis* strains were routinely grown in Brain Heart Infusion (BHI) broth at 37 °C without agitation.

### Bacteriophage lysate preparation

To prepare the host culture, an overnight culture of *B. subtilis* pDG1662 was sub-cultured 1:100 into 20 mL LB. The culture was incubated at 37 °C and 220 rpm for 4 hours until the OD_600nm_ reached 0.2. About 1×10^3^ plaque forming units (PFUs) of bacillus phage SPO1 were added to the culture. The phage-infected culture was incubated at 37 °C and 220 rpm until bacterial cells were lysed and the culture turned clear. The phage lysate was filtered through a 0.2 µm polyethersulfone filter and stored at 4 °C.

Pseudomonas phage Lit1 was similarly propagated on *P. aeruginosa* PAO1:Tn7 empty, and the host was cultured in LB + 10 mM MgSO_4_. Enterococcus phage NPV1 was similarly propagated on *E. faecalis* OG1RF, and the host was cultured in Todd-Hewitt broth (THB) + 10 mM MgSO_4_.

### Strain construction

#### Construction of *B. subtilis* that carry Thoeris systems

The Thoeris cassette with its native promoter from *B. amyloliquefaciens* Y2 [NCBI accession #CP003332, 2071378-2073427 (−)] and *B. cereus* MSX-D12 [AHEQ01000050, 16453-19685 (+)] were synthesized and cloned into the HindIII site on plasmid pDG1662 by GenScript (Plasmid maps included as Supplementary Files 1, 2). The constructed plasmids were propagated in *E. coli* DH5α and selected by 100 µg/mL ampicillin. The plasmids were extracted from *E. coli* DH5a using QIAprep Spin Miniprep Kit (QIAGEN #27104). Then the plasmids were transformed into *B. subtilis* RM125 using a protocol adapted from Young et al.^51^ Briefly, *B. subtilis* RM125 were grown in Medium A (1 g/L yeast extract, 0.2 g/L casamino acids, 0.5% (w/v) glucose, 15 mM (NH_4_)_2_SO_4_, 80 mM K_2_HPO_4_, 44 mM KH_2_PO_4_, 3.9 mM sodium citrate, 0.8 mM MgSO_4_) at 37 °C, 220 rpm until the cessation of logarithmic growth and were allowed to continue growing for another 90 mins. Then *B. subtilis* cells were diluted 10-fold into Medium B (Medium A + 0.5 mM CaCl_2_ + 2.5 mM MgCl_2_) and incubated at 37 °C, 300 rpm for 90 mins. 1 µg of plasmid was added to these now-competent cells and incubated at 37 °C, 220 rpm for 30 mins. Transformed *B. subtilis* cells with the double-crossover insertion at the *amyE* locus were selected based on resistance to 5 µg/mL chloramphenicol and sensitivity to 100 µg/mL spectinomycin.^52^

#### Construction of *B. subtilis* that expresses Y2 ThsB under IPTG induction

Briefly, Y2 ThsB was placed downstream of the P_spank_ promoter (IPTG-inducible) on plasmid pDR110 and then integrated at the *amyE* locus on *B. subtilis* genome. First, Y2 ThsB was amplified by PCR using pDG1662:Y2 Thoeris as the template with primers Y2ThsB_pDR110_F and Y2ThsB_pDR110_R, followed by DpnI digestion and PCR cleanup using QIAquick PCR Purification Kit (QIAGEN #28104). pDR110 was then amplified by PCR using primers pDR110_F and pDR110_R, followed by DpnI digestion and gel purification using QIAquick Gel Extraction Kit (QIAGEN #28704). Y2 ThsB was ligated with pDR110 using NEBuilder® HiFi DNA Assembly Master Mix (New England Biolab #E2621) and electroporated into NEB® 10-beta Electrocompetent *E. coli* (New England Biolab #C3020K), which was selected by 100 µg/mL ampicillin. The sequence of the constructed plasmid was verified by whole plasmid sequencing (Plasmid map included as Supplementary Files 3). The plasmid was extracted from *E. coli* NEB® 10-beta using QIAprep Spin Miniprep Kit before transformation into *B. subtilis* RM125 using the above protocol. Transformed *B. subtilis* cells were selected with resistance to 100 µg/mL spectinomycin. Colonies with double-crossover insertion were verified by colony PCR using primers amyE_F and amyE_R. All PCR reactions were performed using Q5® Hot Start High-Fidelity 2X Master Mix (New England Biolab #M0494).

### Construction of *E. faecalis* that carries the Thoeris system

Briefly, the DS16 Thoeris operon was placed downstream of the P*_bacA_* promoter (constitutively active) on plasmid pLZ12A, and then the P_bacA_*-*DS16 Thoeris cassette was genomically integrated between genes OG1RF_11778 and OG1RF_11779 in *E. faecalis* OG1RF using shuttle vector pWH03.^53^ To construct the pLZ12A vector carrying DS16 Thoeris, the genomic DNA of *E. faecalis* DS16 was extracted from 1 mL of overnight culture using Wizard® Genomic DNA Purification Kit (Promega #A1120). The Thoeris operon from the *E. faecalis* DS16 genome [NCBI accession AJEY01000012.1, 80649-82108 (−)] was amplified by PCR using primers DS16_Thr_F and DS16_Thr_R, followed by PCR cleanup using QIAquick PCR Purification Kit. pLZ12A was amplified by PCR using primers pLZ12A_F and pLZ12A_R, followed by DpnI digestion and PCR cleanup using QIAquick PCR Purification Kit. DS16 Thoeris was ligated with pLZ12A using NEBuilder® HiFi DNA Assembly Master Mix and transformed by heat shock into NEB® 5-alpha Competent *E. coli* (New England Biolab #C2987) and selected by 15 µg/mL chloramphenicol. The sequence of the constructed plasmid was verified by whole plasmid sequencing (Plasmid map included as Supplementary File 4).

To construct the pWH03 vector carrying P_bacA_*-*DS16 Thoeris, the P_bacA_ promoter with DS16 Thoeris operon was amplified by PCR using pLZ12A:DS16Thr as the template with primers pLZ12A_DS16Thr_F and pLZ12A_DS16Thr_R, followed by DpnI digestion and gel purification using QIAquick Gel Extraction Kit. The integration vector pWH03 was amplified by PCR using primers pWH03_F and pWH03_R, followed by DpnI digestion and gel purification using QIAquick Gel Extraction Kit. P_bacA_*-*DS16 Thoeris was ligated with pWH03 using NEBuilder® HiFi DNA Assembly Master Mix and electroporated into NEB® 10-beta Electrocompetent *E. coli* which was selected by 15 µg/mL chloramphenicol. The sequence of the constructed plasmid was verified by whole plasmid sequencing (Plasmid map included as Supplementary File 5). To generate the P_bacA_*-*DS16 Thoeris genomic insertion mutant, pWH03:DS16Thr was electroporated into electro-competent *E. faecalis* OG1RF cells and selected as described previously.^54^ The presence of the pWH03:DS16Thr in OG1RF cells was validated by colony PCR using two pairs of primers: pheS_198F and EF2238_200R, EF2238_827F and DS16ThsA_200R. The single-site integration by homologous recombination at either EF2238 or EF2239 was validated by colony PCR using primer pairs OG1RF11777_60F and DS16ThsA_200R or DS16ThsB_321F and OG1RF11780_94R respectively. The genomic integration of P_bacA_*-*DS16 Thoeris between EF2238 and EF2239 was validated by colony PCR using primers OG1RF11777_60F and OG1RF11780_94R. All PCR reactions were performed using Q5® Hot Start High-Fidelity 2X Master Mix (New England Biolab #M0494).

### Construction of *P. aeruginosa* that carries the Thoeris system

The Thoeris type II locus from Pa MRSN11538 was integrated into the PAO1 genome under regulation of a constitutively active promoter. For chromosomal insertion of Thoeris type II at the Tn7 locus in *P. aeruginosa* PAO1 (PAO1:Thoeris II), the integrating vector pUC18-mini-Tn7T-LAC^55^ carrying the Thoeris type II operon was used along with the transposase-expressing helper plasmid pTNS3. The pUC18-mini-Tn7-Thoeris II vector was used for the creation of the PAO1:Thoeris II strain, and the pUC18-Tn7T-LAC empty vector was used for the creation of PAO1:Tn7 empty strain, which was used as a negative control. For insertion of Thoeris II, its operon was PCR amplified from the MRSN11538 *Pa* strain genomic DNA using primers Ths_Pa_11538_F and Ths_Pa_11538_R. The PCR product was gel purified using Monarch DNA gel extraction kit and inserted into the HindIII/BamHI-cleaved pUC18-Tn7T-LAC vector using NEBuilder HiFi DNA Assembly. Next, a constitutively active promoter followed by ribosome binding site was inserted upstream the Thoeris II locus to allow constitutive expression of ThsB and ThsA genes. To obtain this construct, the plasmid from the previous cloning step was PCR amplified using primers pUC18-Tn7_const_Promoter_F and Tn7_const_Promoter_R, followed by DpnI treatment, gel purification, and self-ligation using NEBuilder HiFi DNA Assembly. The resulting plasmids were used to transform into *E. coli* DH5ɑ. The sequence of the constructed plasmid was verified by whole plasmid sequencing (Plasmid map included as Supplementary File 6). *P.aeruginosa* PAO1 cells were electroporated with the pUC18-mini-Tn7-Thoeris II vector or pUC18-mini-Tn7T-LAC and pTNS3, and the resulting strains were selected on gentamicin-containing plates. Potential integrants were screened by colony PCR. Electrocompetent cell preparations, transformations, integrations, selections, plasmid curing, and FLP-recombinase-mediated marker excision with pFLP were performed as described previously.^55^

### High-throughput screening

A synthetic compound library from ChemBridge was used for the screen. The compounds from the library were prepared as 40 µM in LB + 4% DMSO, and 10 µL of these stock solutions were added into the wells of 384-well plates. An overnight culture of *B. subtilis* Y2 Thoeris was diluted 1:100 into fresh LB media + 5 µg/mL chloramphenicol and incubated at 37 °C, 220 rpm for 2 hours. Then, 20 µL of the freshly grown *B. subtilis* culture was added into each well of 384-well plates, followed by incubation at 37 °C for 1 hour. Then, 10 µL of SPO1 phage in LB (∼1000 PFUs) was added to each well. The plates were incubated at 37 °C in a Biospa8 (Biotek) and the OD_600nm_ in each well was recorded every 1 hour using a Synergy H1 plate reader (Biotek).

The Z-score was calculated by

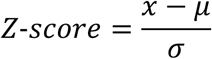

where:

*x* is the OD_600nm_ of the well at 10 hours after phage infection,

*µ* is the mean of OD_600nm_ across the plate at 10 hours after phage infection,

*σ* is the standard deviation of OD_600nm_ across the plate at 10 hours after phage infection.

### Evaluation of Thoeris protection on solid media

The sensitivity of bacteria hosts to phages on solid media was determined by the small drop plaque assay. In brief, 100 μL of an overnight culture of *B. subtilis* host was mixed in 5 mL 55 °C LB + 0.5% agar and poured on top of an LB + 1.5% agar plate. After the top soft agar layer solidified, 5 µL of 1:10 dilutions of SPO1 in LB was dropped on top of the soft agar. After the phage spot dried, the plate was incubated at 37 °C overnight.

For *P. aeruginosa,* 10 mM MgSO_4_ was supplemented into LB media, LB + 0.35% agar was used as top agar. 2.5 µL of 1:10 dilutions of Lit1 phage were dropped on top of the soft agar.

For *E. faecalis*, THB media + 10 mM MgSO_4_ was used for cell growth and phage dilution.

### Evaluation of Thoeris protection in liquid media

The sensitivity of bacteria hosts to phages in liquid media was determined by monitoring the growth curve of bacteria. When testing compounds, compounds were dissolved in DMSO or water depending on their solubility, and 2 µL of these stock solutions were added into wells of a 96-well plate. When not testing compounds, 2 µL of solvent was added into each well. An overnight culture of *B. subtilis* host was diluted 1:100 into fresh LB. Then, 180 µL of the diluted culture of *B. subtilis* was added into wells of 96-well plate. The plate was incubated at 30 °C for 30 mins before 20 µL of SPO1 phage (∼10,000 PFUs) in LB media was added into each well. For no infection control, 20 µL of media was added. The plate was then incubated in a Synergy H1 plate reader (Biotek) at 30 °C, 208 rpm (5 mm orbital shaking) and the growth curve of bacteria was recorded by monitoring OD_600nm_ every 30 min.

For *P. aeruginosa*, the 96-well plate containing compounds was prepared as described above. An overnight culture of *P. aeruginosa* host was diluted 1:100 into fresh LB + 10 mM MgSO_4_. Then, 180 µL of the diluted culture of *P. aeruginosa* was added into wells of 96-well plate. The plate was incubated at 37 °C for 30 mins before 20 µL of Lit1 phage (∼10,000 PFUs) in LB + 10 mM MgSO_4_ was added into each well. For no infection control, 20 µL of media was added. The plate was incubated at 37 °C in a Biospa8 (Biotek) and the OD_600nm_ in each well was recorded every 30 min using a Synergy H1 plate reader (Biotek).

For *E. faecalis*, the 96-well plate containing compounds was prepared as described above. An overnight culture of *E. faecalis* host was diluted 1:1000 into fresh THB + 10 mM MgSO_4_. Then, 180 µL of the diluted culture of *E. faecalis* was added into wells of 96-well plate. The plate was incubated at 37 °C for 1 hour before 20 µL of NPV1 phage (∼1,000 PFUs) in THB + 10 mM MgSO_4_ was added into each well. For no infection control, 20 µL of media was added. The plate was incubated at 37 °C in a Biospa8 (Biotek) and the OD_600nm_ in each well was recorded every 30 min using a Synergy H1 plate reader (Biotek).

The strength of the Thoeris system under compound treatment was calculated by

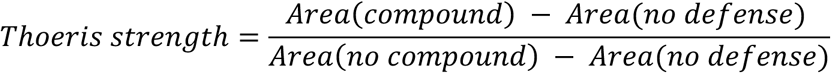

where “area” represents the integrated area under the bacterial lysis curve upon phage infection (**Figure S2a**). The Thoeris strength of the no compound control group was defined as 1, while the Thoeris strength of no defense control was defined as 0.

### Phage reproduction measurement in liquid media

Phage reproduction was evaluated by quantifying the number of phages produced after infecting bacterial hosts. When testing compounds, compounds were dissolved in DMSO or water depending on the solubility and 2 µL of these solutions were added into wells of a 96-well plate. When not testing compounds, 2 µL of solvent (DMSO or water) was added into each well. An overnight culture of *B. subtilis* host was diluted 1:100 into fresh LB. Then, 180 µL of the diluted culture of *B. subtilis* was added into the wells of 96-well plate. The plate was incubated at 30 °C for 30 mins before 20 µL of SPO1 phage (∼10,000 PFUs) in LB media was added into each well. The plate was then incubated in a Synergy H1 plate reader (Biotek) at 30 °C, 208 rpm (5 mm orbital shaking) and the growth curve of bacteria was recorded by monitoring OD_600nm_ every 30 min. After 15 hours of incubation, 200 µL of the infected culture was removed and centrifuged at 16,000 g for 10 mins. The PFUs in the supernatant were quantified on *B. subtilis* without defense using the small drop plaque assay described above.

For *P. aeruginosa*, the 96-well plate containing compounds was prepared as described above. An overnight culture of *P. aeruginosa* host was diluted 1:100 into fresh LB + 10 mM MgSO_4_. Then, 180 µL of the diluted culture of *P. aeruginosa* was added into wells of 96-well plate. The plate was incubated at 37 °C for 30 mins before 20 µL of Lit1 phage (∼10,000 PFUs) in LB + 10 mM MgSO_4_ was added into each well. The plate was incubated at 37 °C in a Biospa8 (Biotek) and the OD_600nm_ in each well was recorded every 30 min using a Synergy H1 plate reader (Biotek). After 15 hours of incubation, 200 µL of the infected culture was removed and centrifuged at 16,000 g for 10 mins. The PFUs in the supernatant were quantified on *P. aeruginosa* without defense using the small drop plaque assay described above.

For *E. faecalis*, the 96-well plate containing compounds was prepared as described above. An overnight culture of *E. faecalis* host was diluted 1:1000 into fresh THB + 10 mM MgSO_4_. Then, 180 µL of the diluted culture of *E. faecalis* was added into wells of 96-well plate. The plate was incubated at 37 °C for 1 hour before 20 µL of NPV1 phage (∼1,000 PFUs) in THB + 10 mM MgSO_4_ was added into each well. The plate was incubated at 37 °C in a Biospa8 (Biotek) and the OD_600nm_ in each well was recorded every 30 min using a Synergy H1 plate reader (Biotek). After 15 hours incubation, 200 µL of the infected culture was removed and centrifuged at 16,000 g for 10 mins. The PFUs in the supernatant were quantified on *E. faecalis* without defense using the small drop plaque assay described above.

### Preparation of phage-infected cell lysate for LC-HRMS analysis

Lysates were prepared as described previously^12^ with minor modifications described below. Overnight cultures of *B. subtilis* P_spank_ or *B. subtilis* P_spank_-Y2 ThsB were diluted 1:100 into 500 mL fresh LB + 100 µg/mL spectinomycin + 1 mM IPTG. When testing inhibitors, 500 µM of the compound was supplemented to the *B. subtilis* P_spank_-Y2 ThsB culture. The diluted cultures were incubated at 30 °C, 220 rpm for 4 hours until OD_600nm_∼0.3. 50 mL of the culture was removed as a t=0 min sample and immediately centrifuged at 10,000 g at 4 °C for 5 mins. The supernatant was discarded, and the cell pellet was stored at −80 °C. Then, 5 mL of SPO1 phage (∼5 ×10^10^ PFUs/mL) was added to the host cells to reach MOI∼10. The infected cell culture was incubated at 30 °C, 220 rpm, and 50 mL of the culture was removed at different time points to be immediately centrifuged at 10,000 g, 4 °C for 5 mins. The supernatant was discarded, and the cell pellet was stored at −80 °C. The cell pellets were thawed at room temperature and resuspended in 600 µL of 100 mM sodium phosphate buffer (pH = 7) + 4 mg/mL lysozyme. After incubation at room temperature for 10 mins, the cells were transferred into 2 ml tubes with Lysing Matrix B (MP Biomedicals #116911050) and lysed using an Omni Bead Ruptor 12 for 2 × 40 s at 6 m/s with a dwell time of 4 mins in-between. After lysis, the tubes were centrifuged at 15,000 g at 4 °C for 10 mins. Then, 400 µL of each supernatant were transferred to Amicon Ultra-0.5 Centrifugal Filter Units 3 kDa (EMD Millipore #UFC500396) and centrifuged for 45 mins at 14,000 g 4 °C. The filtrate was collected, and 10 µl of each were used for LC-MS analysis.

### LC-MS analysis of His-ADPR, inhibitor-ADPR, and NAD^+^ in the phage infected-cell lysate

The liquid chromatography analysis was performed on ACQUITY UPLC I-Class PLUS System using a Luna Omega 5 μm Polar C18 100 Å column (250×4.6 mm). The mobile phase A was water + 0.1 % (v/v) formic acid and the mobile phase B was acetonitrile + 0.1 % (v/v) formic acid. The flow rate was kept at 0.7 mL·min^-1^ and the gradient was as follows: 0% B (0–10 min), increase to 2.5% B (10–15 min), increase to 5% B (15–16 min), hold 5% B (16–26 min), increase to 95% B (26–27 min), hold 95% B (27–37 min), decrease to 0% B (37–38 min), hold 0% B (38– 48 min). High-resolution electrospray ionization (HR-ESI) mass spectra with collision-induced dissociation (CID) MS/MS were obtained using a Waters Synapt G2S Quadrupole Time-of-Flight (QTOF). The instrument was operated at negative ionization mode. The MS spectra were obtained on the Time-of-Flight analyzer with a scan range of 300–800 Da and analyzed using MassLynx 4.1 software. The *m/z* of interest was filtered through Quadrupole, subjected to CID (energy ramp 34–44 V), and analyzed on the Time-of-Flight analyzer with a scan range of 50 – 750 Da.

### IP6C-ADPR and His-ADPR pull-down with BaY2ThsB and BaY2ThsA^Macro^ proteins

A 0.4 L culture of *E. coli* TOP10 cells harboring the vector pBAD_DelTM-ThsA-TwinStrep_ThsB-His^12^ was induced at OD600 0.6 with 0.2% L-arabinose, IP6C was added to the culture to the final concentration of 1.25 mM and cells were grown overnight at 16 °C. Control cells were induced without the addition of IP6C. The cells were harvested by centrifugation and re-suspended in (1) the Strep-Wash buffer (100 mM Tris-HCl pH 8.0, 150 mM NaCl, 1 mM EDTA, 2 mM phenylmethylsulfonyl fluoride (PMSF)) for BaY2ThsA^Macro^ purification, or (2) His-Wash buffer for (10 mM sodium phosphate pH 8.0, 150 mM NaCl, 0.01% Tween-20, 2 mM PMSF) for BaY2ThsB purification, or (3) 20 mM Tris-HCl pH 8.0 buffer for lysate control, and lysed by sonication. After removing debris by centrifugation, the supernatants were mixed with MagStrep® Strep-Tactin®XT beads (IBA, cat no. 2-5090-002) for BaY2ThsA purification or Dynabeads™ His-Tag Isolation and Pulldown beads (Invitrogen™, cat no. 10103D) for BaY2ThsB purification, accordingly. Protein purification was performed according to manufacturers’ protocols. Purified protein was denatured for 5 mins at 98°C and centrifuged for 15 mins at 16,000 *g*. Control lysate was transferred to Amicon Ultra-0.5 Centrifugal Filter Unit 3 kDa and centrifuged for 30 mins at 4°C, 12,000 *g*. Resulting supernatant and filtrate were analyzed by LC-MS (see below).

### In-vitro formation of IP6C-ADPR catalyzed by BaY2ThsB and ArThsB

To produce IP6C-ADPR in vitro, reactions containing 1 mM NAD^+^, 3 mM L-Histidine, 10 mM IP6C and 100 μM ThsB were prepared in the reaction buffer containing 10 mM Na-HEPES (pH 7.5 at 25°C), 150 mM NaCl and 5 mM MgCl_2_ and incubated for 1 day at 25°C and later 6 days at 37°C. Samples were heat-denatured for 5 min at 98°C, centrifuged for 15 min at 16,000 g and the resulting supernatants were analyzed by LC-MS (see below).

### LC-MS analysis of the in vitro formed and pulled-down molecules

LC-MS analysis was carried out on 1290 Infinity HPLC system (Agilent Technologies) coupled to a 6520 Accurate Mass Q-TOF LC–MS mass analyzer (Agilent Technologies) with an electrospray ion source. HPLC was carried out on a Supelco Discovery HS C18 column at a temperature of 30 °C. Chromatography was carried out at a 0.3 mL‧min^−1^ flow rate using a linear mobile phase gradient over 30 min 0.02% formic acid in water to 0.02% formic acid in acetonitrile. MS was carried out using gas at 300 °C, 10 L‧min^−1^ gas flow, 2,500 V capillary voltage, 150 V fragmentator voltage. Data acquisition and analysis were carried out using QTOF Acquisition Software (B.02.01 SP1) and MassHunter (vB.05.00, Agilent Technologies) software.

### Production and purification of BaY2ThsA^Macro^

The BaY2ThsA gene were synthesized as a gBlock (Integrated DNA Technologies). BaY2ThsA^Macro^ (residues 83-328) was amplified by polymerase chain reaction and cloned into the pET28b vector using Gibson Assembly reaction.^56^ The resulting construct was verified by sequencing. BaY2ThsA^Macro^ in the pET28B vector [C-terminal twin Strep-tag] was produced in *E. coli* BL21 (DE3) cells, using the autoinduction method, and purified to homogeneity, using a combination of Strep-tag affinity chromatography and SEC. Briefly, the cells were grown at 37°C, until an optical density at 600 nm of 0.6 to 0.8 was reached. The temperature was then reduced to 20°C, and the cells were grown overnight for approximately 16 hours. The cells were harvested by centrifugation at 5000*g* at 4°C for 15 min and stored at −80°C. The cell pellets were resuspended in 2 to 3 ml of lysis/wash buffer [20 mM Tris (pH 8.0), 1 M NaCl, 0.5 mM TCEP, 0.1% TRITON X-100 and 5% v/v glycerol] per gram of cells. The resuspended cells were lysed using a sonicator and clarified by centrifugation (15,000*g* for 30 min). The clarified lysate was applied to a Strep-Tactin XT 4flow cartridge (IBA) pre-equilibrated with 10 CVs of the lysis/wash buffer at a rate of 0.5 ml/min. The column was washed with 10 CVs of the wash buffer, followed by elution of bound proteins using lysis/wash buffer supplemented with 50 mM D-biotin. The elution fractions were analyzed by SDS-PAGE, and the fractions containing the protein of interest were pooled and further purified on a S200 HiLoad 26/600 column pre-equilibrated with gel filtration buffer [20 mM Tris (pH 8.0), 0.5 M NaCl, 0.2 mM TCEP and 5% v/v glycerol]. The peak fractions were analyzed by SDS-PAGE, and the fractions containing BaY2ThsA^Macro^ were pooled and concentrated to final concentrations of approximately 1.6 mg/ml, flash-frozen as 10-μl aliquots in liquid nitrogen, and stored at −80°C.

### NMR Spectroscopy for enzymatic reaction, STD-NMR, and IP6C-ADPR standard

NMR samples were prepared in a total volume of 200 μL consisting of 175 µL HBS buffer (50 mM HEPES, 150 mM NaCl, pH 7.5), 20 µL D_2_O, and 5 µL DMSO-d6. Each sample was subsequently transferred to a 3 mm Bruker NMR tube rated for 600 MHz data acquisition. All ^1^H NMR spectra were acquired with a Bruker AVANCE NEO 600 MHz NMR spectrometer equipped with quadruple resonance QCIF CryoProbe at 298 K. To suppress resonance from H_2_O, a water-suppression pulse program (P3919GP), using a 3-9-19 pulse-sequence with gradients,^57, 58^ was implemented to acquire spectra with an acquisition delay of 2 s and 32 scans per sample. For enzymatic reaction, ^1^H spectra were recorded at multiple time-points depending on instrument availability. The pulse-sequence STDDIFFGP19.3, in-built within the TopSpin^TM^ program (Bruker), was employed to acquire STD-NMR spectra.^59^ The on-resonance irradiation was set close to protein resonances at 0.8 ppm, whereas the off-resonance irradiation was set far away from any protein or ligand resonances at 300 ppm. A relaxation delay of 4 s was used, out of which a saturation time of 3 s was used to irradiate the protein with a train of 50 ms Gaussian shaped pulses. The number of scans was 256. All spectra were processed by TopSpin™ 4 (Bruker) and Mnova 14 (Mestrelab Research).

### Synthesis and purification of IP6C-ADPR standard

IP6C-ADPR standard was produced via TIR domain catalysed base-exchange using NAD^+^ and IP6C as substrates. A 10 mL sample of 0.5 µM His_6_-tagged *Bacillus subtilis* SpbK,^60^ 5 mM IP6C, and 10 mM NAD^+^ in HBS buffer (50 mM HEPES, 150 mM NaCl, pH 7.5) with 2.5% DMSO was incubated at room temperature and reaction progress was monitored intermittently by ^1^H NMR over time. To stop the reaction, the His_6_-tagged enzyme was removed by incubating the mixture with 200 μL of HisPur™ Ni-NTA resin for 30-60 min. The resin was subsequently removed by centrifugation at 500 x *g* for 1 min and the supernatant was subjected to HPLC-based separation to purify the base-exchange products. A Shimadzu Prominence HPLC equipped with a Synergi™ 4 µm Hydro-RP 80 Å column was used for separation. The mobile phase consisted of phase A (0.04 % (v/v) TFA in water) and phase B (0.04 % (v/v) TFA in acetonitrile). Different gradients, flow rates, and run times were applied depending on prior optimization with individual reaction mixtures. Product peaks were confirmed by comparison with individual chromatograms of NAD^+^, nicotinamide, ADPR, and IP6C. Fractions corresponding to the IP6C-ADPR peak were collected, concentrated, lyophilized, and stored at −20°C. NMR characterizations of IP6C-ADPR were performed by the aforementioned NMR spectrometer and the detailed peak assignment can be found in **Table S4**.

### In vivo phage therapy experiment in mouse model

This protocol was adapted from a previous study.^61^ The Animal Research Ethics Committee of the Army Medical University reviewed, approved and supervised the protocols for animal research (permit number: AMUWEC20240067). The mice were purchased from Hunan SJA Laboratory Animal Company and housed under specific pathogen-free conditions; the housing environment had controlled temperature (20–26 °C), humidity (40–70%) and lighting conditions (12 h light and 12 h dark cycle), and no animal was excluded from the analyses.

For the toxicity test of the compounds, 80 μl of compound IP6C or Id5C (12.5 mg/mL) were injected intraperitoneally at 0h, 12h and 24h. Each group included 7 mice, which were observed for 7 days. After 7 days post-infection, mice that survived the initial challenge were euthanized.

For the phage therapy model experiment, the PAO1:Thoeris II strain was cultured in LB at 37 °C until the early stationary phase. Cells were then collected and resuspended in PBS to OD600 of 0.6. A volume of 50 μl (∼3 × 10^6^ CFUs) of bacteria suspension was intraperitoneally inoculated into 7-week-old BALB/c female mice. Immediately following the bacteria infection, a volume of 50 μl of Lit1 phage (3×10^7^ or 3×10^8^ PFUs) was inoculated intraperitoneally on the other side, followed by the intraperitoneal injection of 80 μl of compound IP6C or Id5C (12.5 mg/mL) at 0h, 12h and 24h after the inoculation of phage. Each group included 7 mice, which were observed for 7 days. After 7 days post-infection, mice that survived the initial challenge were euthanized.

### Synthesis of compound 27-28

**Scheme S4.1.**
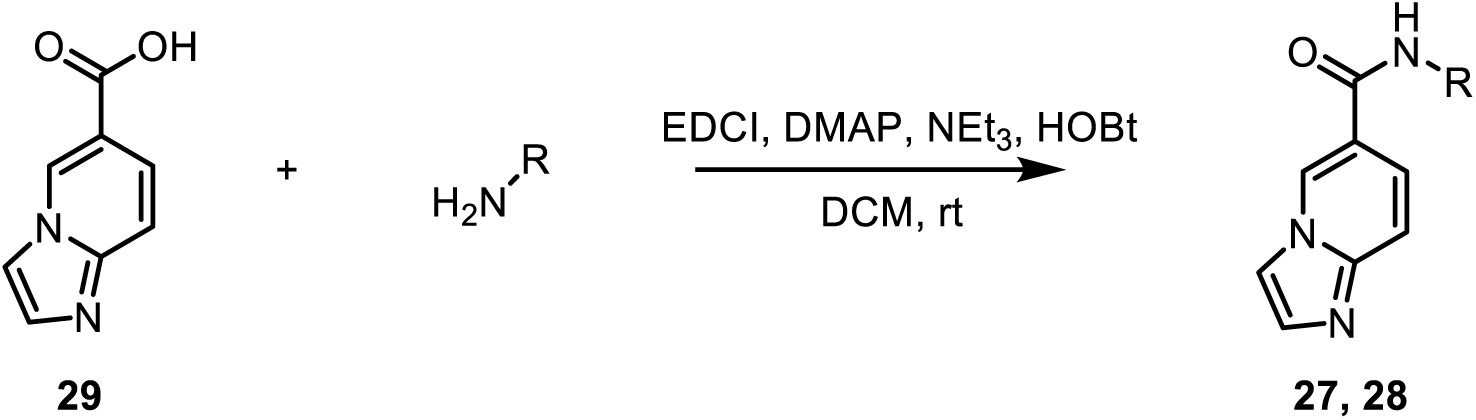
Synthesis of compound 27-28.

The ^1^H NMR spectra were obtained on a Varian 600 MHz Inova NMR spectrometer using Varian/Agilent VnmrJ and Linux workstations. All the spectra were analyzed using MestReNova 14.2.0-26256 software.

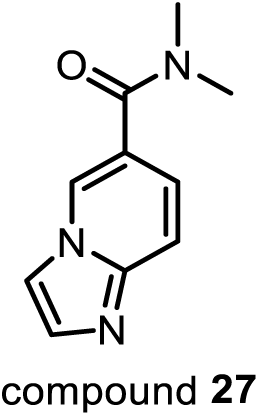

*N,N-dimethylimidazo[1,2-a]pyridine-6-carboxamide (**27**)*. Compound **29** (50 mg, 308 µmol), EDCI (65.0 mg, 339 µmol), DMAP (7.5 mg, 62 µmol), NEt_3_ (152 mg, 1.5 mmol), and HOBt (45.8 mg, 339 µmol) were added to 20 mL CH_2_Cl_2_. The mixture was stirred on ice for 1 hour, followed by addition of dimethylamine (27.8 mg, 616 µmol). The reaction mixture was then stirred at room temperature for 18 hours before being washed with 60 mL of saturated Na_2_CO_3_. The organic layer was dried over anhydrous Na_2_SO_4_ and concentrated in vacuo. The crude product was purified by silica gel (CH_2_Cl_2_:MeOH = 20:1) to give compound **27** (20.8 mg, 35% yield) as a pale white solid. ^1^H NMR (600 MHz, CD_2_Cl_2_) δ 8.35 (t, *J* = 1.3 Hz, 1H), 7.66 (t, *J* = 1.0 Hz, 1H), 7.63 (d, *J* = 1.3 Hz, 1H), 7.57 (dt, *J* = 9.2, 0.9 Hz, 1H), 7.20 (dd, *J* = 9.2, 1.7 Hz, 1H), 3.05 (s, 6H). Spectrum is shown in **Figure S12**.

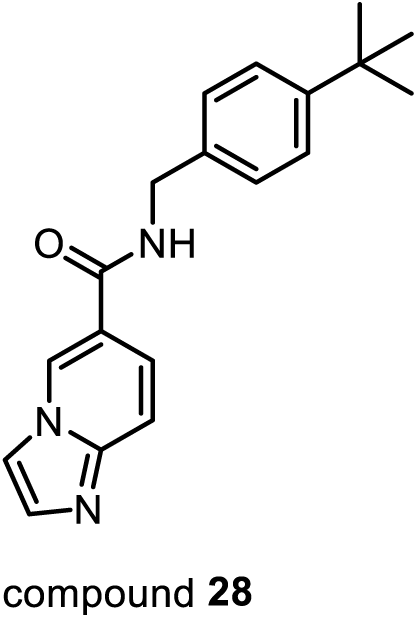

*N-(4-(tert-butyl)benzyl)imidazo[1,2-a]pyridine-6-carboxamide (**28**)*. Compound **29** (50 mg, 308 µmol), EDCI (65.0 mg, 339 µmol), DMAP (7.5 mg, 62 µmol), NEt_3_ (152 mg, 1.5 mmol), and HOBt (45.8 mg, 339 µmol) were added to 20 mL CH_2_Cl_2_. The mixture was stirred on ice for 1 hour, followed by addition of 4-*tert*-butylbenzylamine (100.6 mg, 616 µmol). The reaction mixture was then stirred at room temperature for 18 hours before washed with 60 mL of saturated Na_2_CO_3_. The organic layer was dried over anhydrous Na_2_SO_4_ and concentrated in vacuo. The crude product was purified by silica gel (CH_2_Cl_2_:MeOH = 20:1) to give compound **28** (54.0 mg, 57% yield) as a pale white solid. ^1^H NMR (600 MHz, CD_2_Cl_2_) δ 8.83 (t, *J* = 1.4 Hz, 1H), 7.72 (t, *J* = 5.8 Hz, 1H), 7.56 (dd, *J* = 12.2, 1.6 Hz, 2H), 7.51 (dd, *J* = 9.4, 1.8 Hz, 1H), 7.41 (d, *J* = 9.4 Hz, 1H), 7.35 (d, *J* = 8.3 Hz, 2H), 7.27 (d, *J* = 8.2 Hz, 2H), 4.58 (d, *J* = 5.7 Hz, 2H), 1.31 (s, 9H). Spectrum is shown in **Figure S13**.

## Data Availability

Any requests for data should be addressed to the corresponding author.

## Acknowledgments

We thank the Bacillus Genomic Stock Center (Ohio State University) for providing bacteria and phages. We thank Tamim Mosaiab and Veronika Masic for support with NMR experiments and protein production. We thank Audrone Ruksenaite (Vilnius University) for conducting the MS experiments. The research was supported by a research starter grant from the American Society of Pharmacognosy to J.P.G., a National Science Foundation CAREER award (IOS-2143636) to J.P.G., and a Camille Dreyfus Teacher-Scholar Award (TC-24-028) to J.P.G. Research support was also provided by the National Health and Medical Research Council (Investigator Grant 1196590 to T.V.), the Australian Research Council (Future Fellowship FT200100572 to T.V.), a Discovery Early Career Researcher Award (DE250101258 to Y.S), the National Institutes of Health (R01AI141479 to B.A.D.), the National Key Research and Development Program of China (2021YFA0911200 to S.L), the Research Council of Lithuania (LMTLT) agreement No S-MIP-24-84 (to G.T.), and the Kleberg Foundation (to J.B.D.). Z.Z. was supported in part by the John R. and Wendy L. Kindig Fellowship. The Laboratory for Biological Mass Spectrometry was supported by the Indiana University Precision Health Initiative. The 600 MHz spectrometer of the Indiana University NMR facility was supported by NSF grant CHE-1920026, and the Prodigy probe was purchased in part with support from the Indiana Clinical and Translational Sciences Institute, funded in part by NIH Award TL1TR002531.

## Supplementary tables and figures

**Figure S1.**
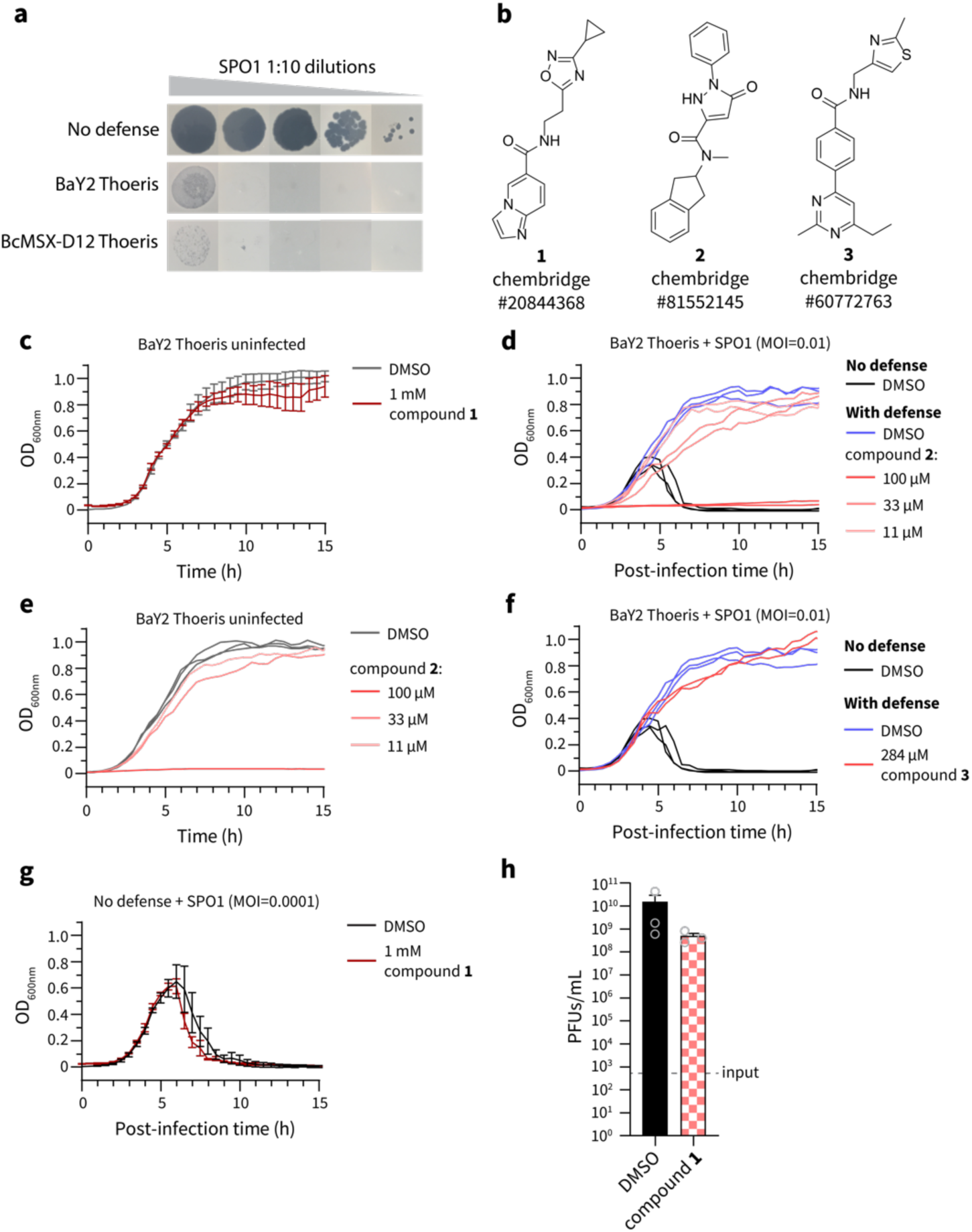
Compound 1 is a specific inhibitor of the BaY2 Thoeris system. (a) BaY2 Thoeris and BcMSX-D12 Thoeris protect *B. subtilis* from SPO1 phage infection on solid media. (b) Chemical structures of compounds **1** – **3**. (c) Compound **1** has no growth effect on *B. subtilis* cells containing BaY2 Thoeris in the absence of phages. Data are represented as the average ± SEM from three independent biological replicates. (d, e) Compound **2** inhibits growth of *B. subtilis* cells containing BaY2 Thoeris, which is independent of SPO1 phage infection, suggesting that compound **2** is an antibacterial agent instead of a Thoeris inhibitor. Data from two independent biological replicates are shown here. (f) The anti-Thoeris effect of compound **3** failed to reproduce after retesting. Data from two independent biological replicates are shown here. (g) Compound **1** does not improve phage infectivity on *B. subtilis* without any defense. Data are represented as the average ± SEM from three independent biological replicates. (h) Phage reproduction of SPO1 on *B. subtilis* without any defense, quantified by measuring the plaque forming units (PFUs) after 15 hours post-infection. Input indicates the initial PFUs in the culture. 1 mM of compound **1** was tested. Data are represented as the average ± SEM from three independent biological replicates. Individual replicates are represented by grey circles.

**Figure S2.**
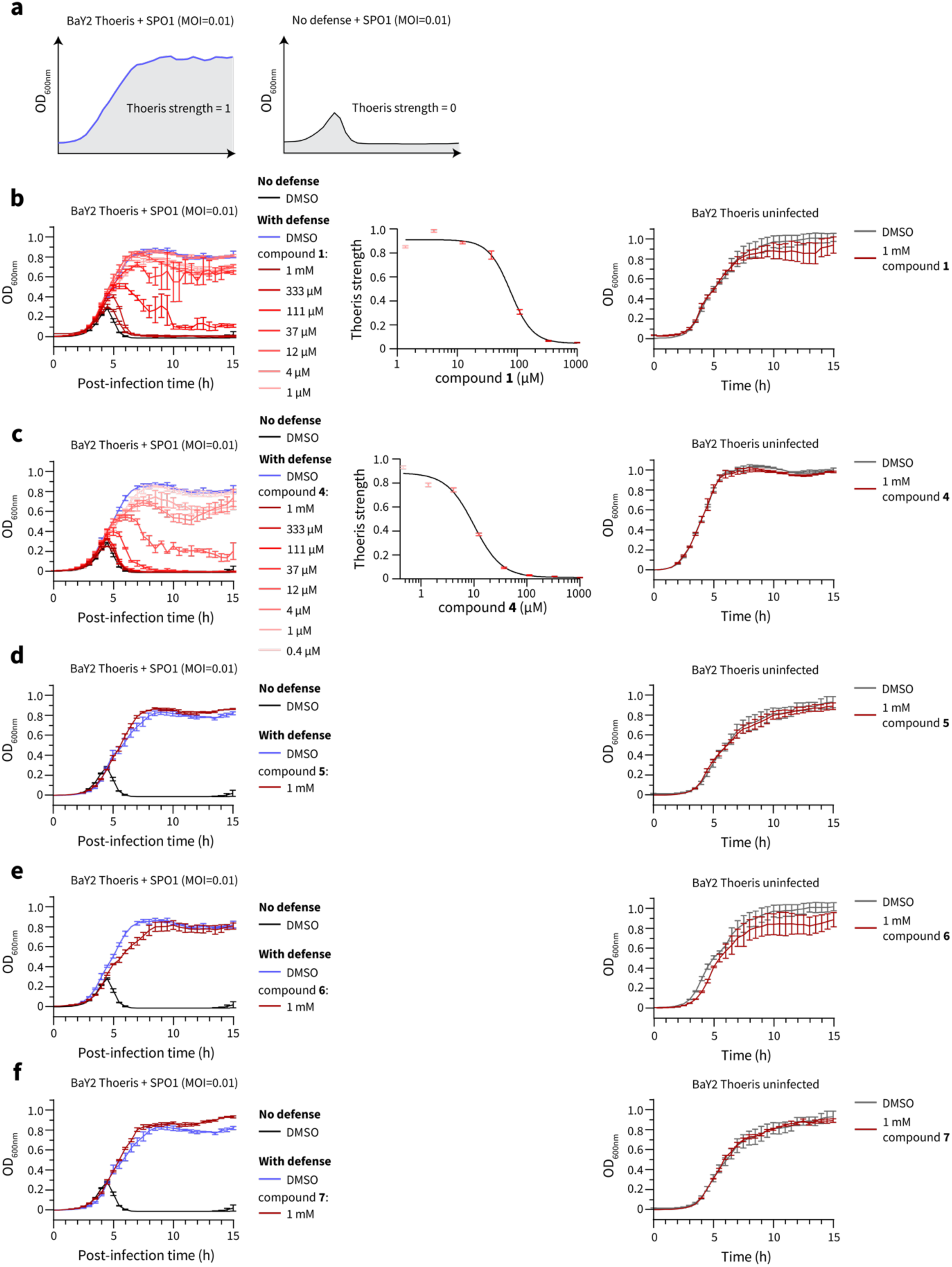
Anti-Thoeris effects and dose-response curves of compounds 1, 4 – 7. (a) Thoeris strength is defined as the area under the lysis curve of *B. subtilis* cells infected by SPO1 phages. The Thoeris strength of *B. subtilis* cells containing BaY2 Thoeris and *B. subtilis* cells without any defense are defined as 1 and 0 respectively. (b – f) The anti-Thoeris effect (left panel), dose-response curve (middle panel, if the compound is active in inhibiting BaY2 Thoeris), and the growth effect (right panel) of compound **1** (b), compounds **4** – **7** (c – f). Data are represented as the average ± SEM from three independent biological replicates.

**Figure S3.**
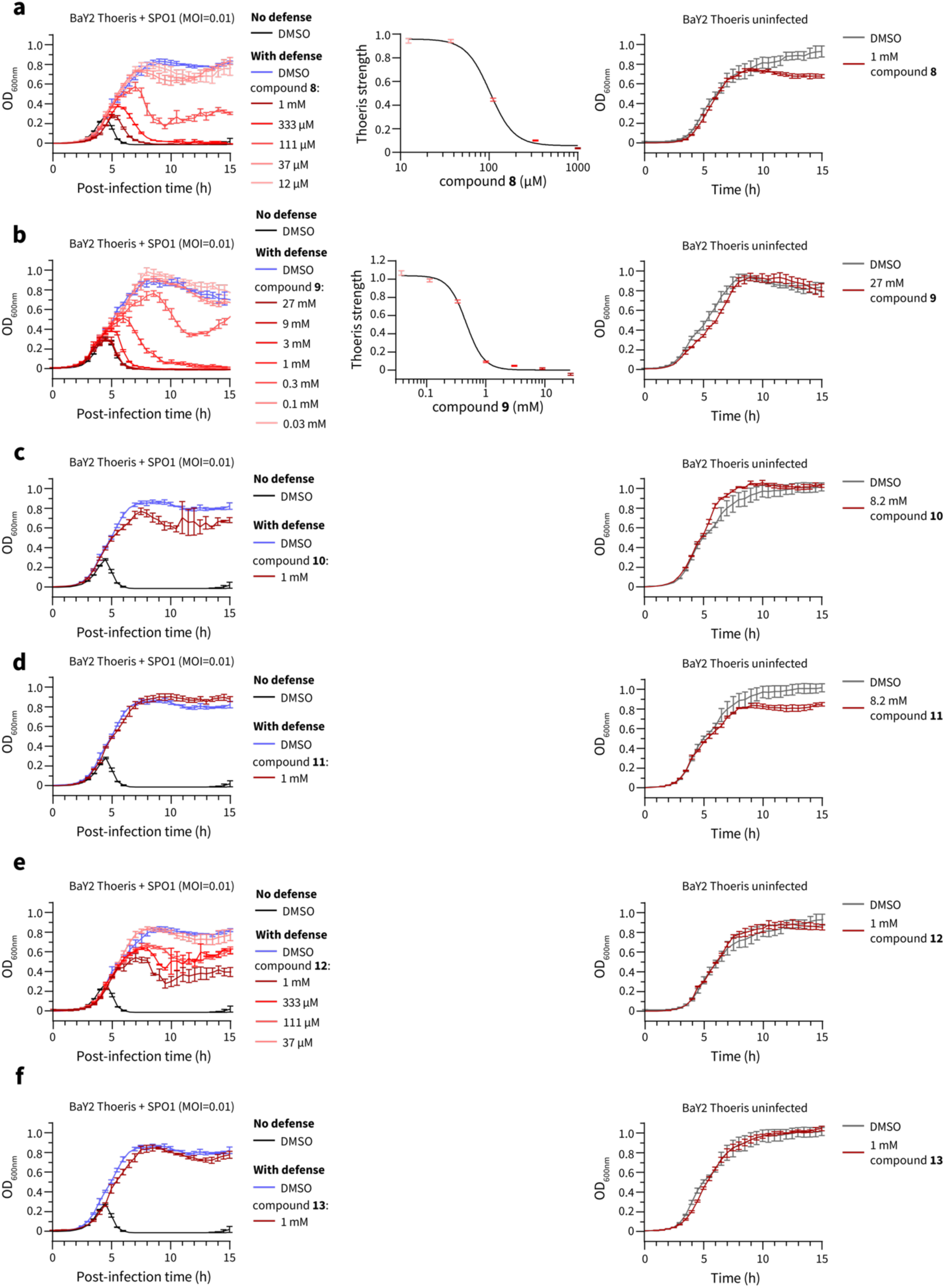
Anti-Thoeris effects and dose-response curves of compounds 8 – 13. The anti-Thoeris effect (left panel), dose-response curve (middle panel, if the compound is active in inhibiting BaY2 Thoeris), and the growth effect (right panel) of compounds **8** – **13** (a – f). Data are represented as the average ± SEM from three independent biological replicates..

**Figure S4.**
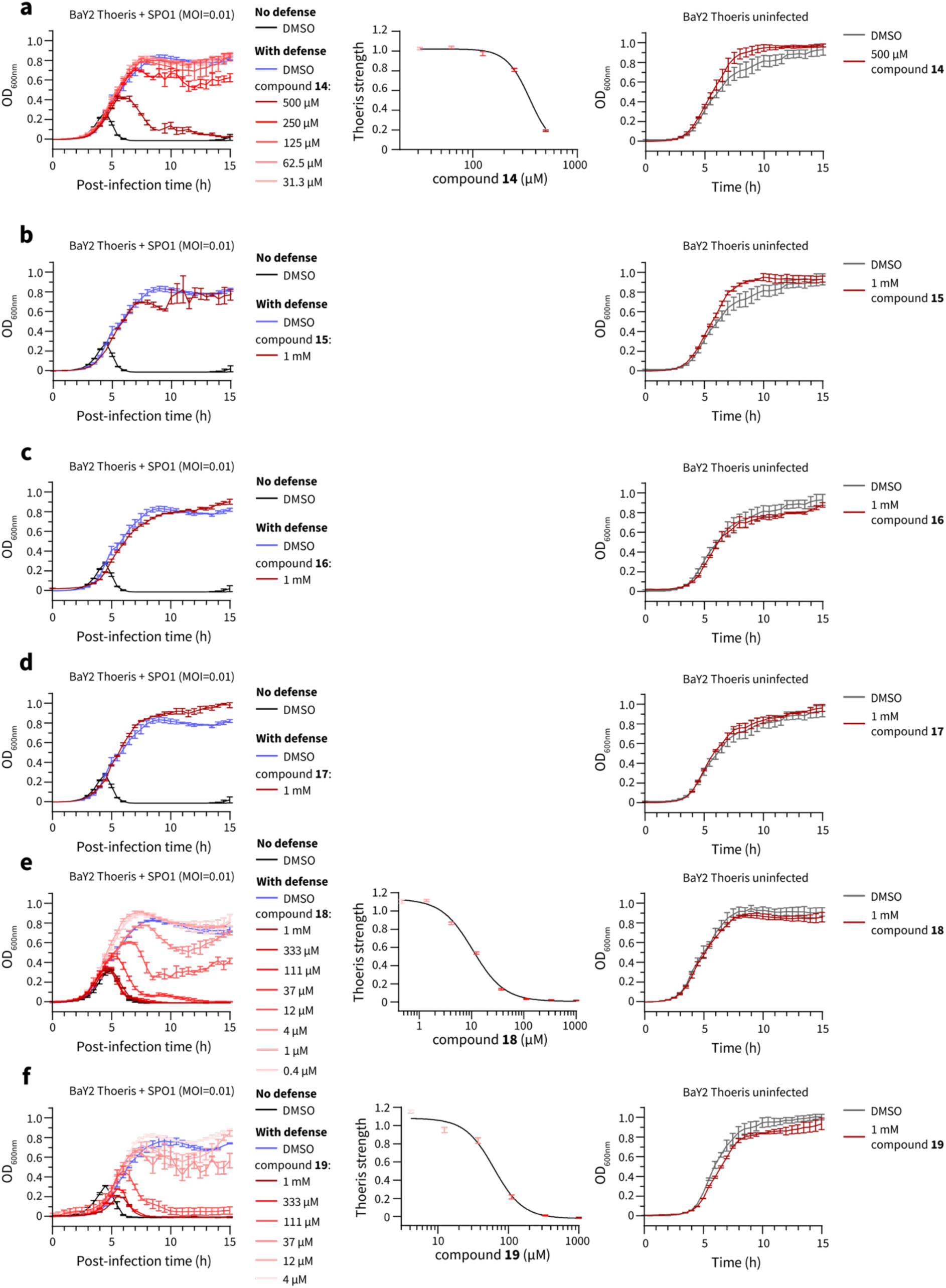
Anti-Thoeris effects and dose-response curves of compounds 14 – 19. The anti-Thoeris effect (left panel), dose-response curve (middle panel, if the compound is active in inhibiting BaY2 Thoeris), and the growth effect (right panel) of compounds **14** – **19** (a – f). Data are represented as the average ± SEM from three independent biological replicates.

**Figure S5.**
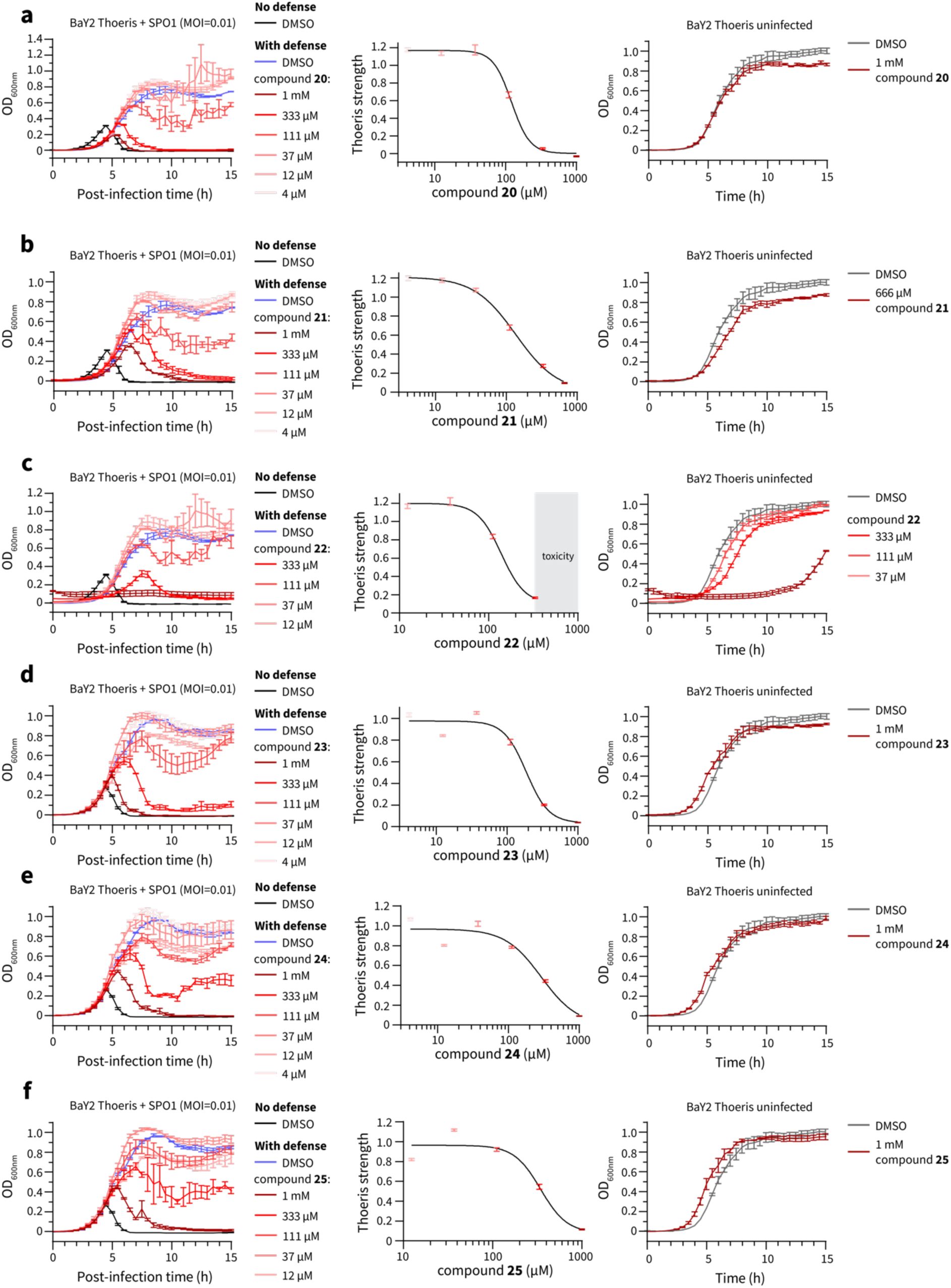
Anti-Thoeris effects and dose-response curves of compounds 20 – 25. The anti-Thoeris effect (left panel), dose-response curve (middle panel, if the compound is active in inhibiting BaY2 Thoeris), and the growth effect (right panel) of compounds **20** – **25** (a – f). Data are represented as the average ± SEM from three independent biological replicates.

**Figure S6.**
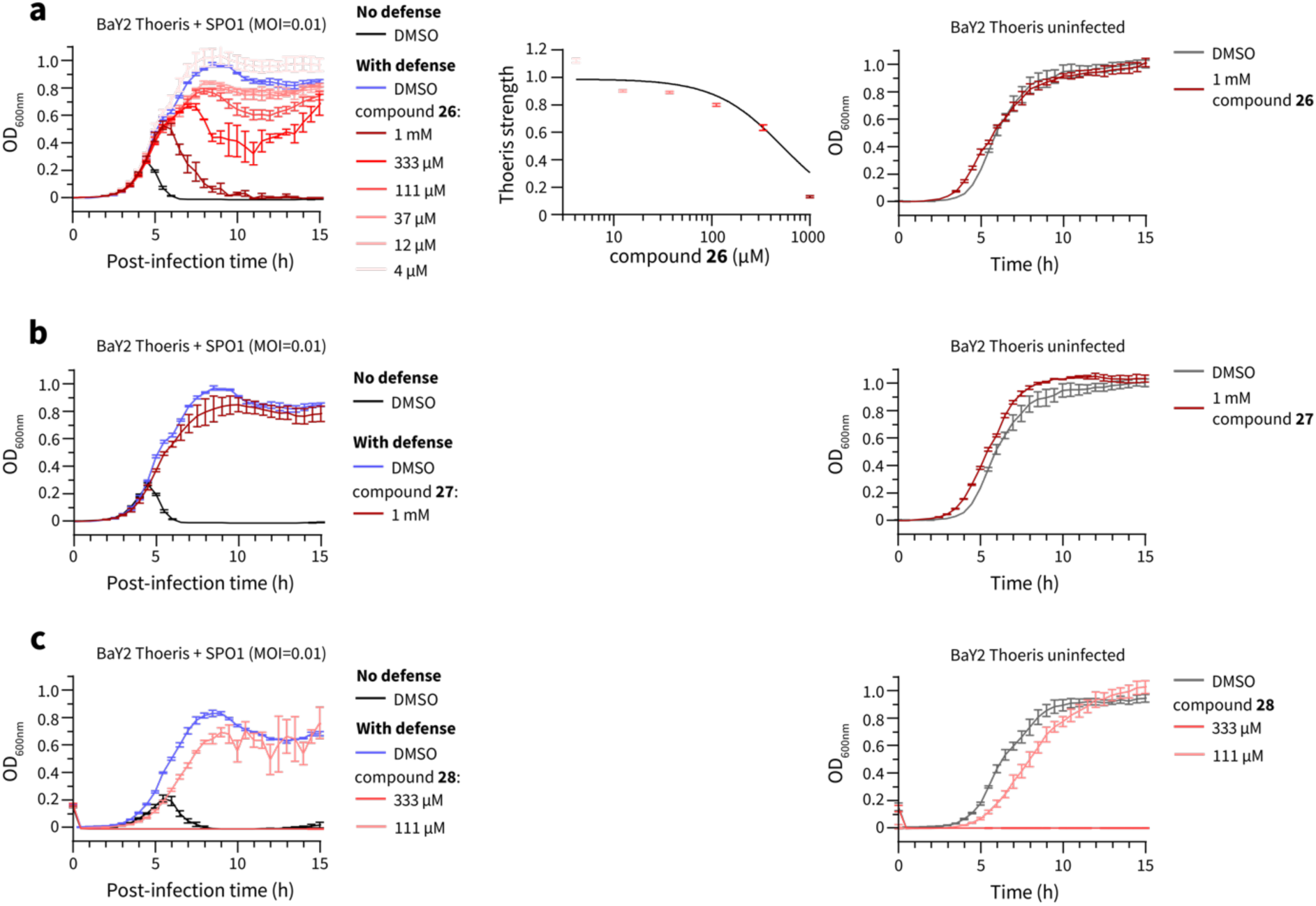
Anti-Thoeris effects and dose-response curves of compounds 26 – 28. The anti-Thoeris effect (left panel), dose-response curve (middle panel, if the compound is active in inhibiting BaY2 Thoeris), and the growth effect (right panel) of compounds **26** – **28** (a – c). Data are represented as the average ± SEM from three independent biological replicates.

**Figure S7.**
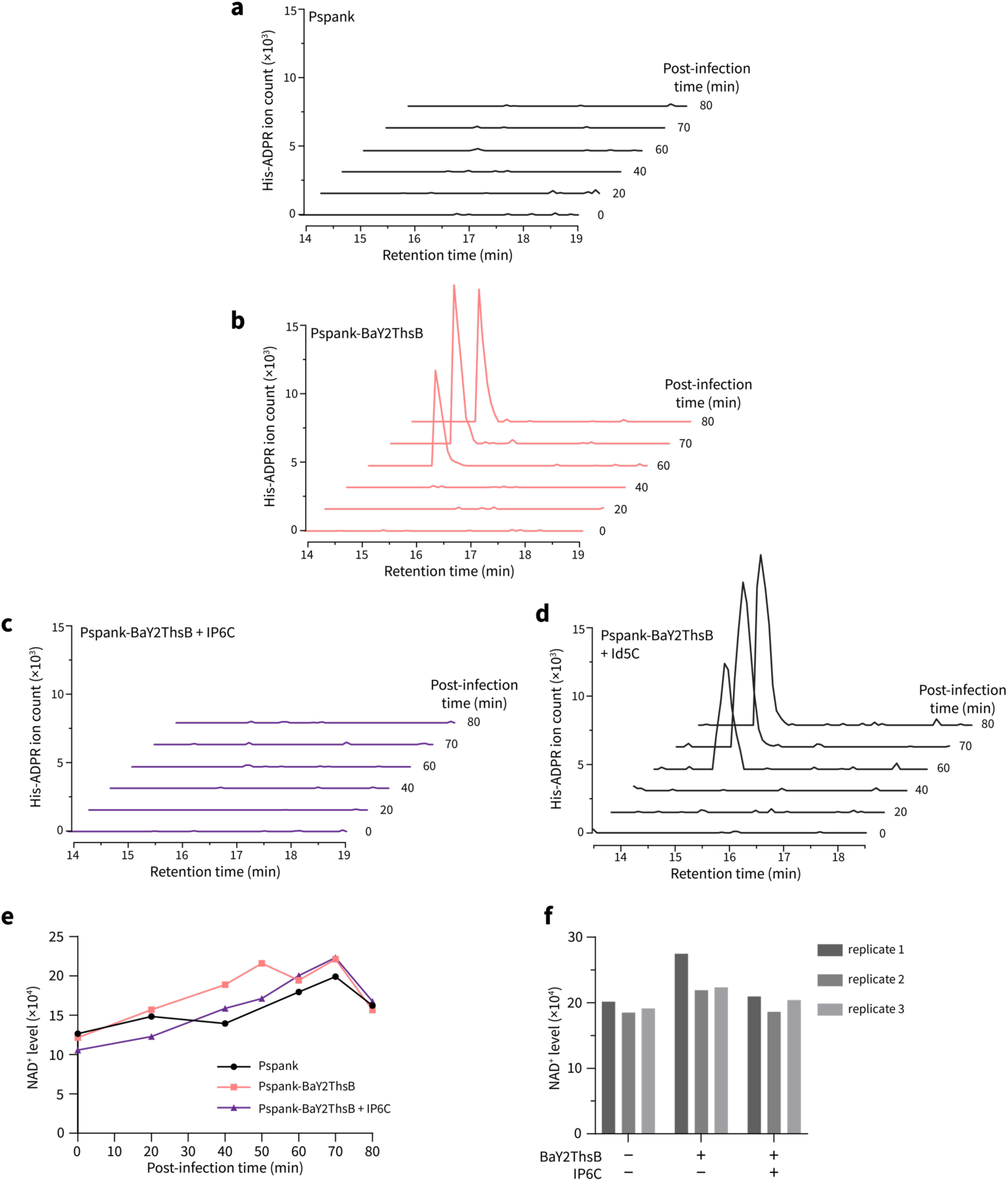
IP6C inhibits His-ADPR signal production by ThsB. (a – d) EICs displaying intracellular His-ADPR levels upon SPO1 phage infection in *B. subtilis* Pspank (a), *B. subtilis* Pspank-BaY2ThsB (b), *B. subtilis* Pspank-BaY2ThsB + 500 µM of IP6C (c), and *B. subtilis* Pspank-BaY2ThsB + 500 µM of Id5C (d). (e) Intracellular NAD^+^ levels reported as the peak area under the extracted ion chromatograms (EICs) of NAD^+^ at each time point after SPO1 infection. 500 µM of IP6C (**4**) was tested and DMSO was used as the negative control. (f) Triplicate measurement of NAD^+^ levels after 80 mins post-infection. 500 µM of IP6C was tested and DMSO was used as the negative control.

**Figure S8.**
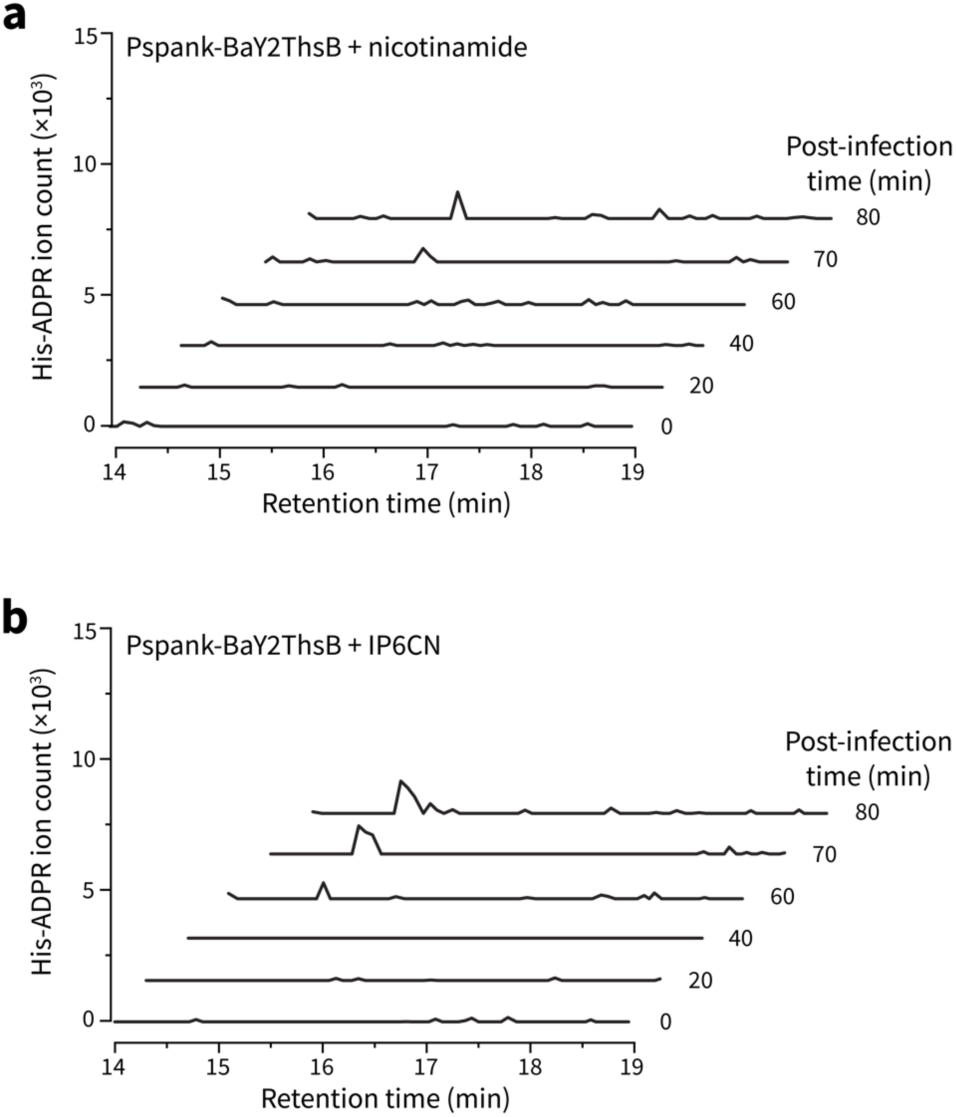
IP6CN (19) and nicotinamide (9) inhibit His-ADPR signal production. EICs displaying intracellular His-ADPR levels upon SPO1 phage infection in *B. subtilis* Pspank-BaY2ThsB in the presence of 8.2 mM nicotinamide (a) or 500 µM of IP6CN (b).

**Figure S9.**
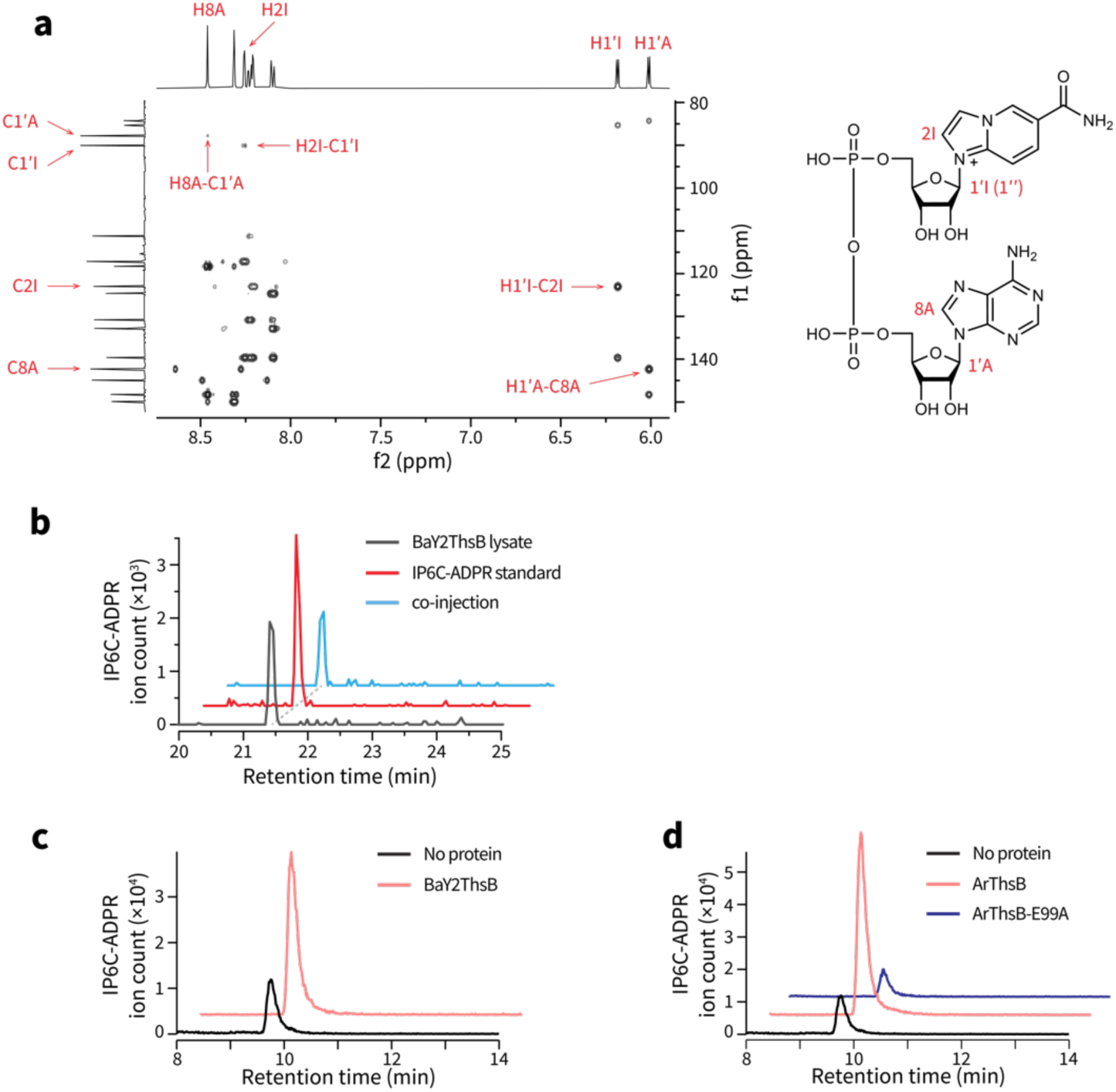
IP6C is converted into IP6C-ADPR by ThsB. (a) Expansion of ^1^H-^13^C HMBC spectrum showing correlations through glycosidic linkages for the IP6C-ADPR standard. (b) Co-injection experiment shows that the IP6C-ADPR made by cells containing BaY2ThsB is the same as the IP6C-ADPR standard. (c) EIC of IP6C-ADPR in the BaY2ThsB reaction mixture. NAD^+^, histidine, and IP6C were used as substrates. (d) EIC of IP6C-ADPR in the ArThsB (BaY2ThsB homolog) reaction mixture. NAD^+^, histidine, and IP6C were used as substrates. E99A mutation abolished the catalytic activity of ArThsB

**Figure S10.**
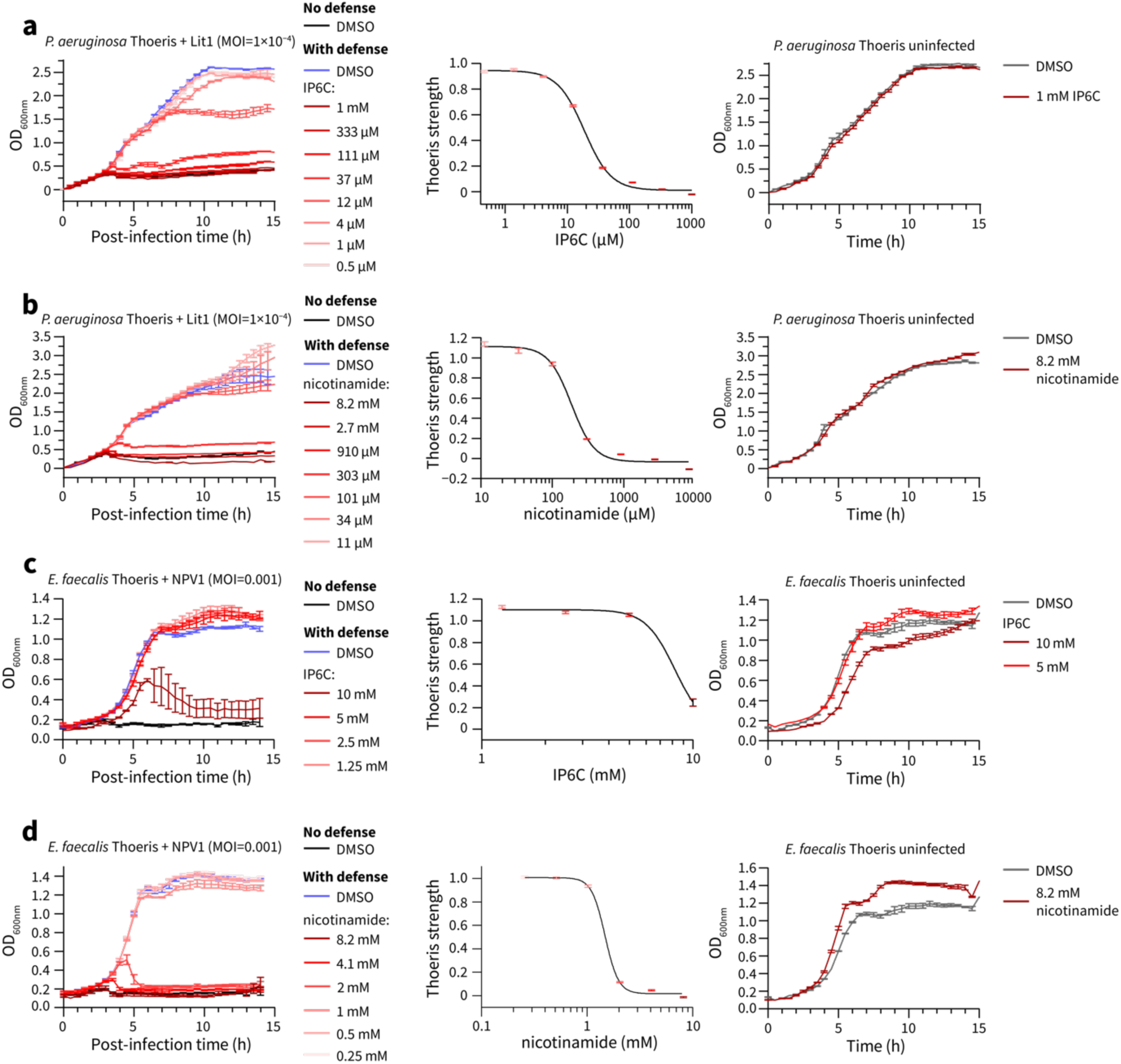
Anti-Thoeris effects and dose-response curves of compound IP6C (4) and nicotinamide (9) in opportunistic pathogens. (a, b) The anti-Thoeris effect (left panel), dose-response curve (middle panel), and the growth effect (right panel) of IP6C (a) and nicotinamide (b) on *P. aeruginosa* Thoeris II. (c, d) The anti-Thoeris effect (left panel), dose-response curve (middle panel), and the growth effect (right panel) of IP6C (c) and nicotinamide (d) on *E. faecalis* Thoeris II. Data are represented as the average ± SEM from three independent biological replicates.

**Figure S11.**
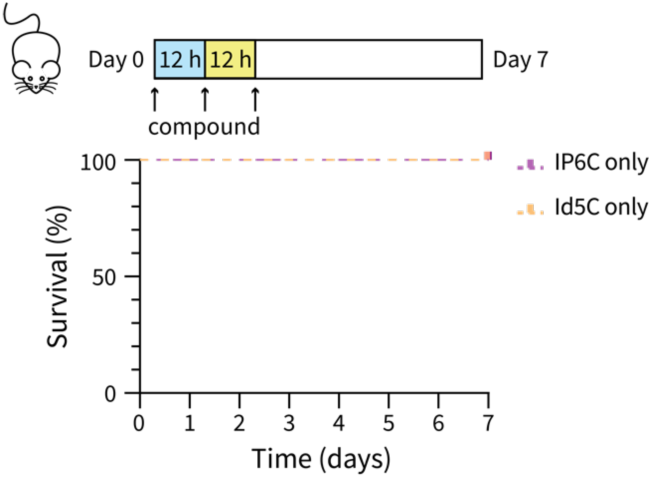
Inhibitors are non-toxic in mice. Survival of 7-week-old BALB/c mice (n = 5) injected with three doses of compounds IP6C and Id5C (each 50 mg/kg) every 12 hours.

**Figure S12.**
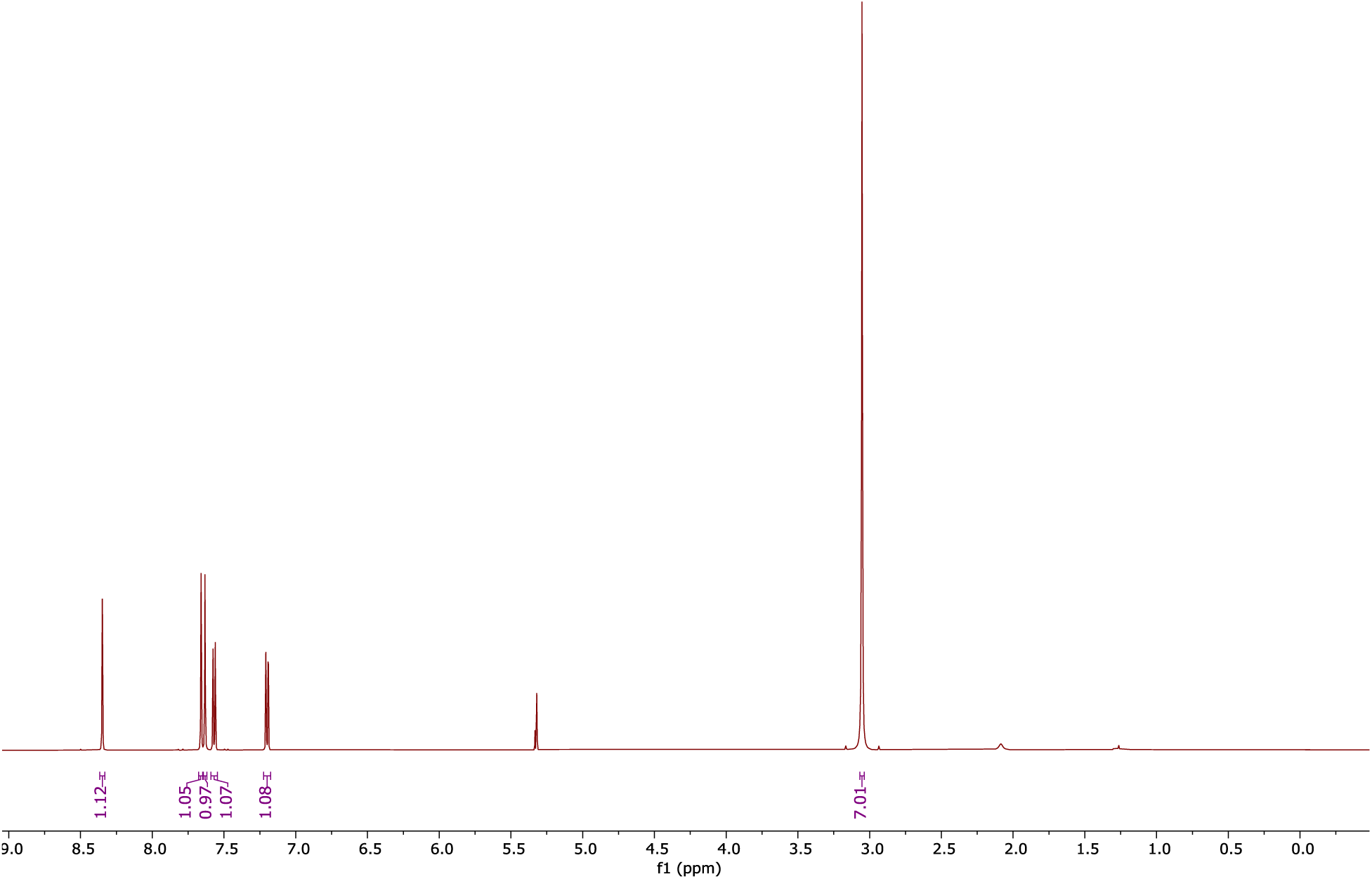
^1^H NMR spectrum (600 MHz) of compound 27 in CD2Cl2.

**Figure S13.**
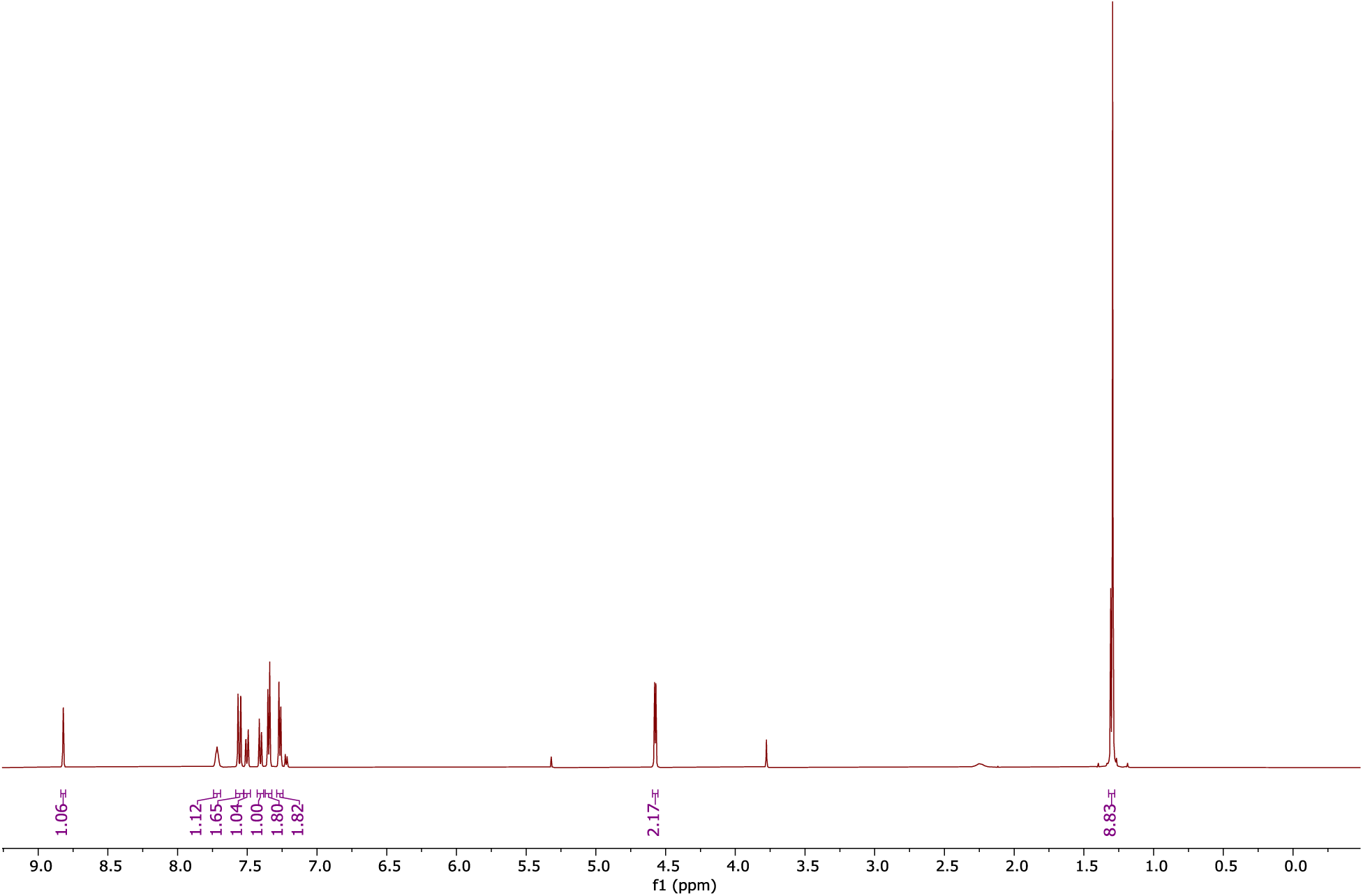
^1^H NMR spectrum (600 MHz) of compound 28 in CD2Cl2.

---

**Table S1.** Strains and bacteriophages used in this study.

| Bacterial strains | Description | Source |
| --- | --- | --- |
| *B. subtilis* RM125 | Wild type strain | Bacillus Genetic Stock Center #1A253 |
| *B. subtilis* pDG1662 | RM125 with pDG1662 inserted at the *amyE* locus on the genome. No Thoeris control. | This study |
| *B. subtilis* Y2 Thoeris | RM125 with pDG1662:Y2 Thoeris inserted at the *amyE* locus on the genome | This study |
| *B. subtilis* MSX-D12 Thoeris | RM125 with pDG1662:MSX-D12 Thoeris inserted at the *amyE* locus on the genome | This study |
| *B. subtilis* Pspank | RM125 with pDR110 inserted at the *amyE* locus on the genome. P<sub>spank</sub> promoter only control. | This study |
| *B. subtilis* Pspank-Y2 ThsB | RM125 with pDR110:Y2 ThsB inserted at the *amyE* locus on the genome. ThsB expression is controlled by an IPTG-inducible promoter (P<sub>spank</sub>) | This study |
| *P. aeruginosa* PAO1:Tn7 empty | PAO1 with empty mini Tn7 vector inserted at *attTn7* site in the genome. No Thoeris control. | This study |
| *P. aeruginosa* PAO1:Thoeris II | PAO1 with Pa MRSN11538 Thoeris II locus inserted at *attTn7* site in the genome | This study |
| *E. faecalis* OG1RF | Wild type strain. No Thoeris control | Lab collection |
| *E. faecalis* DS16 | Clinical isolate carrying Thoeris II system | Lab collection |
| *E. faecalis* GA4 | OG1RF with P<sub>bacA</sub>-DS16 Thoeris II system inserted between OG1RF_11778 and OG1RF_11779 on the genome | This study |
| **Bacteriophages** |  |  |
| SPO1 | *Bacillus* phage | Bacillus Genetic Stock Center #1P4 |
| **Lit1** | *Pseudomonas* phage | Lab collection |
| **NPV1** | *Enterococcus* phage | Lab collection<sup>1</sup> |
| **Plasmids** |  |  |
| **pDG1662** | Empty vector that integrates at *amyE* locus on *Bacillus subtilis* genome | Bacillus Genetic Stock Center #ECE113 |
| **pDG1662:Y2 Thoreris** | Vector carrying Thoreris cassette from *B. amyloliquefaciens* Y2 that integrates at *amyE* locus on *Bacillus subtilis* genome | This study |
| **pDG1662:MSX-D12 Thoreris** | Vector carrying Thoreris cassette from *B. cereus* MSX-D12 that integrates at *amyE* locus on *Bacillus subtilis* genome | This study |
| **pDR110** | Empty vector carrying P<sub>spank</sub> promoter (IPTG-inducible) that integrates at *amyE* locus on *Bacillus subtilis* genome. | Bacillus Genetic Stock Center #ECE311 |
| **pDR110:Y2ThsB** | Vector carrying Y2 ThsB which is downstream of P<sub>spank</sub> promoter that integrates at *amyE* locus on *Bacillus subtilis* genome. | This study |
| **pLZ12A** | Empty vector carries *P<sub>bacA</sub>* promoter for constitutive expression | Lab collection<sup>2</sup> |
| **pLZ12A:DS16Thr** | Vector carries Thoreris operon from *E. faecalis* DS16 which is downstream of *P<sub>bacA</sub>* promoter for constitutive expression | This study |
| **pWH03** | Empty vector that integrates between OG1RF_11778 and OG1RF_11779 on *E. faecalis* OG1RF genome | Lab collection<sup>3</sup> |
| **pWH03:DS16Thr** | Vector carrying *P<sub>bacA</sub>*-DS16 Thoreris that integrates between OG1RF_11778 and OG1RF_11779 on *E. faecalis* OG1RF genome | This study |
| **pUC18- mini-Tn7-Thoreris II** | Vector used for integration of Thoreris II locus in PAO1 genome at *attTn7* site to create PAO1:Thoreris II strain | Lab collection |
| **pUC18- Tn7T-LAC**<br>**empty** | Vector used for creation PAO1: Tn7 empty<br>strain (negative control strain) | Lab collection |

**Table S2.** Chemical used in this study.

| **Chemicals** | **Source** | **Identifier** |
| --- | --- | --- |
| **LB broth** | VWR | Cat#90003-350 |
| **Brain Heart Infusion** | Fisher | Cat#DF0037-17-8 |
| **Todd-Hewitt broth** | Fisher | Cat#DF0492-17-6 |
| **Dehydrated Agar** | Fisher | Cat#DF0140-07-4 |
| **Yeast extract** | Fisher | Cat#DF0127-07-1 |
| **Casamino acids** | Fisher | Cat#DF0231-17-2 |
| **Glucose** | Sigma Aldrich | Cat#G7528 |
| **(NH<sub>4</sub>)<sub>2</sub>SO<sub>4</sub>** | Sigma Aldrich | Cat#A4418 |
| **MgSO<sub>4</sub>** | Sigma Aldrich | Cat#M3634 |
| **Spectinomycin** | Sigma Aldrich | Cat#S9007 |
| **Ampicillin** | Sigma Aldrich | Cat#A9518 |
| **Chloramphenicol** | Sigma Aldrich | Cat#C0378 |
| **DMSO** | Sigma Aldrich | Cat#D8418 |
| **IPTG** | Gold Biotechnology | Cat#I2481C50 |
| **Formic acid (HPLC)** | VWR | Cat#PI85178 |
| **K<sub>2</sub>HPO<sub>4</sub>** | Sigma Aldrich | Cat#S5136 |
| **KH<sub>2</sub>PO<sub>4</sub>** | Sigma Aldrich | Cat#S3139 |
| **CaCl<sub>2</sub>** | Sigma Aldrich | Cat#C7902 |
| **MgCl<sub>2</sub>** | Sigma Aldrich | Cat#63068 |
| **Sodium citrate** | Sigma Aldrich | Cat# C8532 |
| **lysozyme** | Research Products International | Cat#L38100 |
| **DpnI** | New England Biolab | Cat# R0176 |
| **Compound 1** | ChemBridge | Cat#20844368 |
| **Compound 2** | ChemBridge | Cat#81552145 |
| **Compound 3** | ChemBridge | Cat#60772763 |
| **Compound 4** | Combi-Blocks | Cat#QJ-2580 |
| **Compound 5** | ChemBridge | Cat#4041017 |
| **Compound 6** | Sigma Aldrich | Cat#72340 |
| **Compound 7** | Ambeed | Cat#A154365 |
| **Compound 8** | Ambeed | Cat#A226558 |
| **Compound 9** | Ambeed | Cat#A877416 |
| **Compound 10** | AstaTech | Cat#O11834 |
| **Compound 11** | LabNetwork | Cat#EC15116-87-P1 |
| **Compound 12** | Combi-Blocks | Cat#SS-2248 |
| **Compound 13** | Ambeed | Cat#A175137 |
| **Compound 14** | LabNetwork | Cat#EC15116-81-P1 |
| **Compound 15** | LabNetwork | Cat#EC15116-88-P1 |
| **Compound 16** | 1Click Chemistry | Cat#4C70276 |
| **Compound 17** | Ambeed | Cat#A525013 |
| **Compound 18** | Combi-Blocks | Cat#QC-9665 |
| **Compound 19** | Combi-Blocks | Cat#QB-9909 |
| **Compound 20** | Combi-Blocks | Cat#QB-1389 |
| **Compound 21** | Combi-Blocks | Cat#HI-1255 |
| **Compound 22** | Combi-Blocks | Cat#ST-7023 |
| **Compound 23** | Combi-Blocks | Cat#QY-1211 |
| **Compound 24** | Combi-Blocks | Cat#HI-1376 |
| **Compound 25** | Combi-Blocks | Cat#SS-5810 |
| **Compound 26** | Combi-Blocks | Cat#QA-3929 |
| **Compound 29** | Combi-Blocks | Cat#HI-1240 |
| **EDCI** | Sigma Aldrich | Cat#E7750 |
| **DMAP** | Sigma Aldrich | Cat#8510550025 |
| **HOBt** | Sigma Aldrich | Cat#157260 |
| **NEt<sub>3</sub>** | Sigma Aldrich | Cat#TX1200 |
| **Dimethylamine** | Fisher | Cat#AAH27261AE |
| **4-tert-Butylbenzylamine** | Sigma Aldrich | Cat#631280 |

**Table S3.**
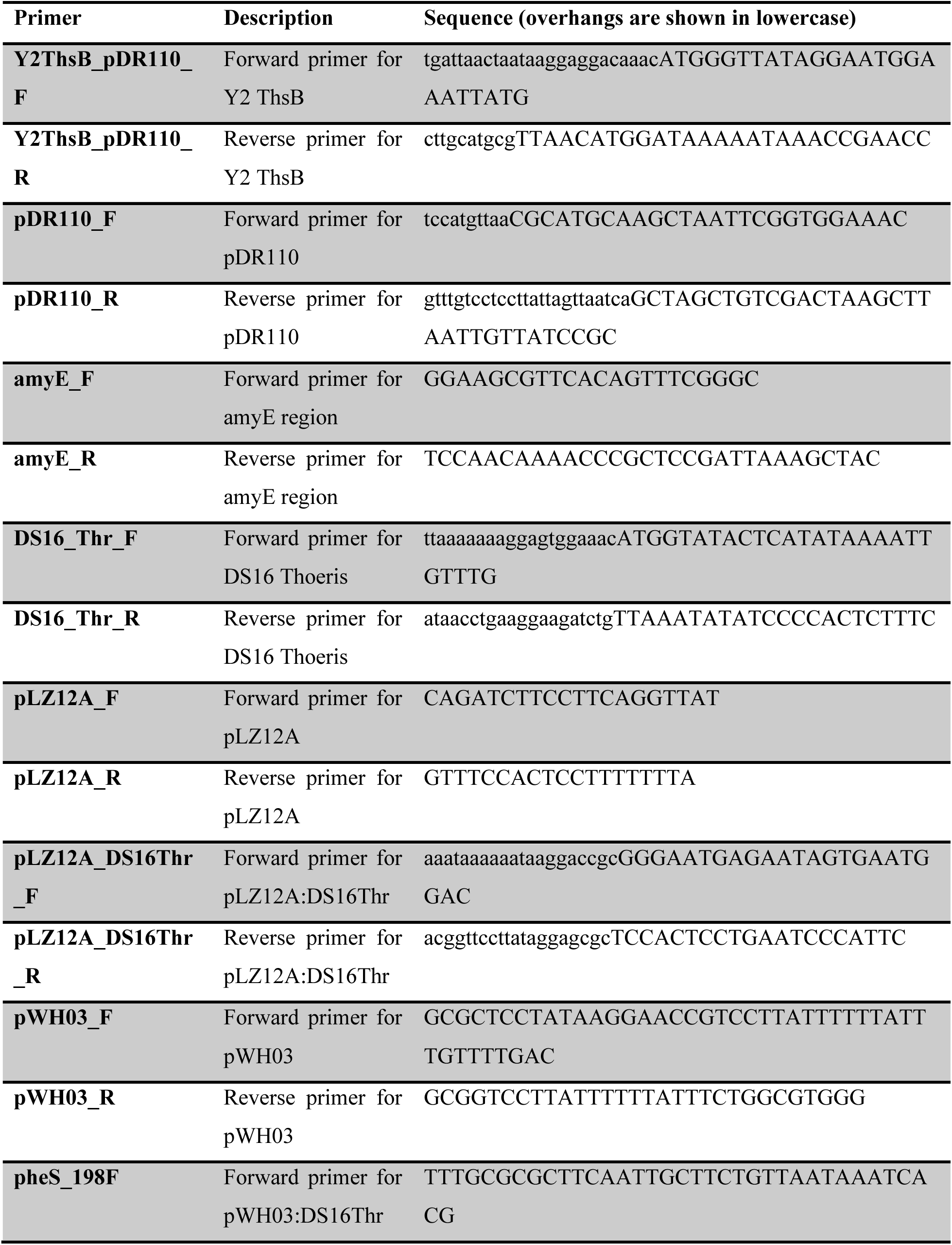
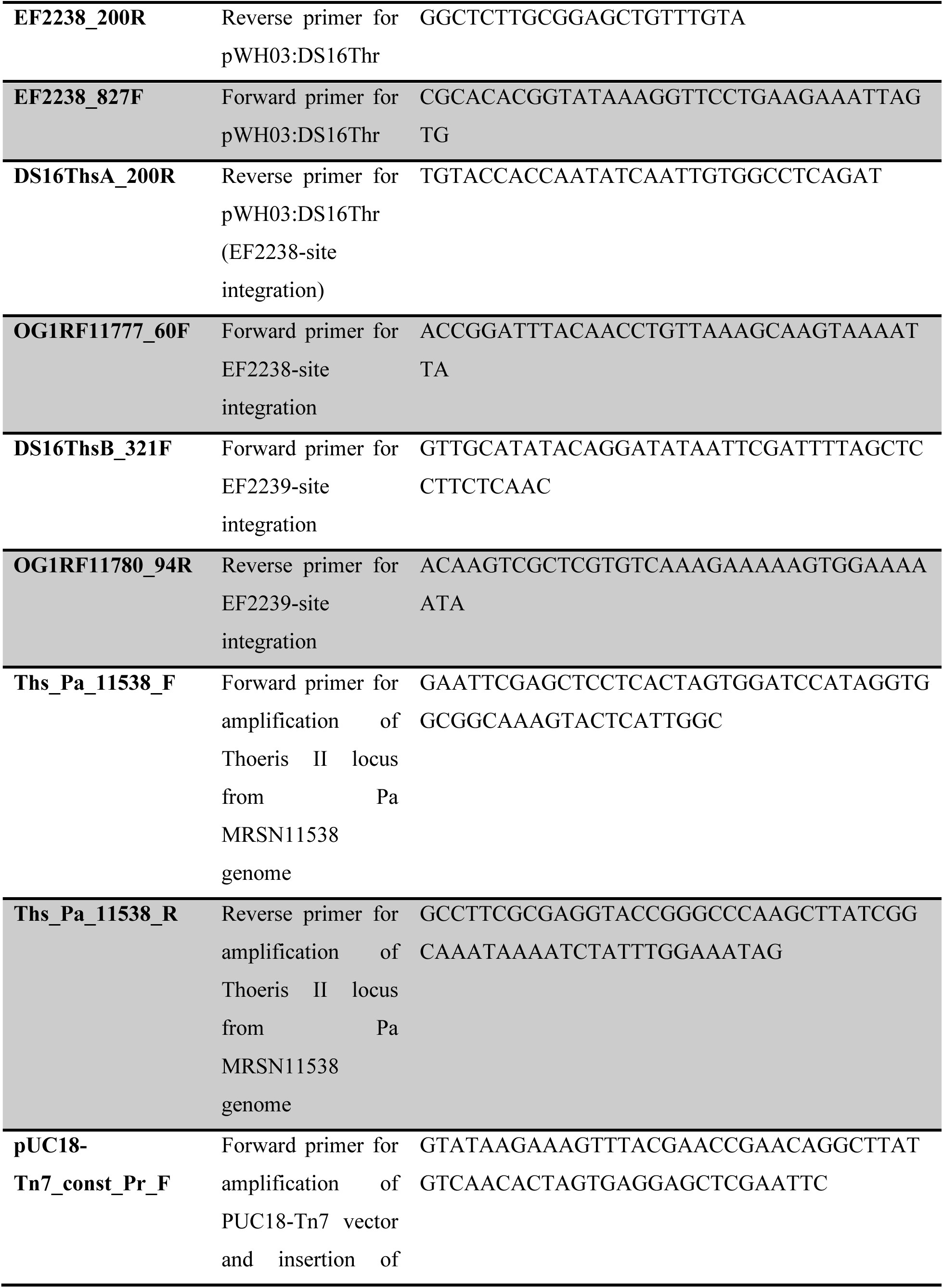
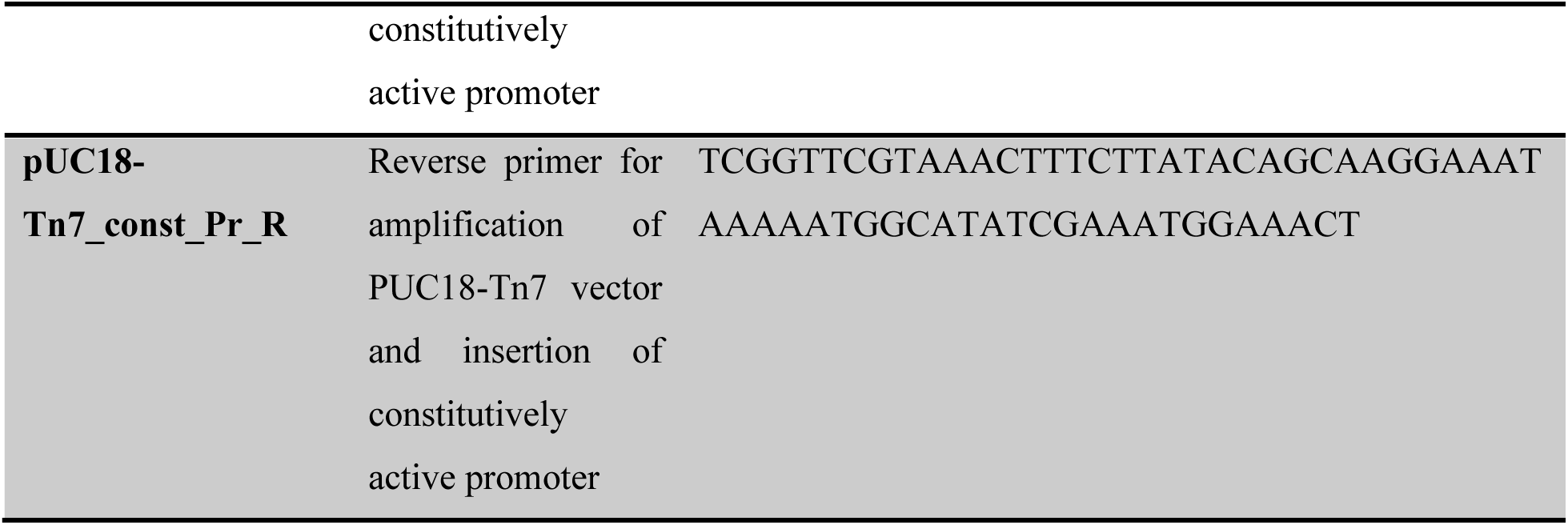
Primers used in this study.

**Table S4.** Assignments of NMR peaks for IP6C-ADPR standard.

| Position | <sup>1</sup>H (ppm, J) | <sup>1</sup>H-<sup>1</sup>H COSY | <sup>13</sup>C (ppm) | <sup>1</sup>H-<sup>13</sup>C HMBC |
| --- | --- | --- | --- | --- |
| 2A | 8.31, s |  | 144.9 | C4A, C6A, C5A |
| 4A |  |  | 148.2 |  |
| 5A |  |  | 118.3 |  |
| 6A |  |  | 149.9 |  |
| 8A | 8.46, s |  | 142.3 | C4A, C6A, C5A, **C1'A** |
| 1'A | 6.01, d (6.0 Hz) | H2'A | 87.8 | **C4A, C8A**, C4'A, C2'A, C3'A |
| 2'A | 4.65, m | H1'A, H3'A | 74.6 | C1'A, C4'A, C3'A |
| 3'A | 4.44, m | H2'A, H4'A | 70.4/70.6 | C1'A, C4'A, C5'A |
| 4'A | 4.30, broad | H3'A, H5'A | 84.3 | C1'A, C2'A, C3'A, C5'A |
| 5'A | 4.19/4.14, broad | H4'A | 65.2 | C4'A, C3'A |
| 2I | 8.26 | H3I | 123.0 | C9I, C3I, **C1'I** |
| 3I | 8.21 | H2I | 117.2 | C9I, C2I |
| 5I | 9.14 |  | 130.8 | C9I, C7I, C6I, C3I, C8I |
| 6I |  |  | 124.6 |  |
| 7I | 8.23 | H8I | 132.8 | C9I, C7I |
| 8I | 8.10 | H7I | 111.2 | C9I, C7I, C5I, C6I |
| 9I |  |  | 139.6 |  |
| 1'I | 6.19, broad | H2'I | 90.1 | **C9I, C2I**, C4'I, C2'I, C3'I |
| 2'I | 4.57, m | H1'I, H3'I | 75.2 | C1'I, C4'I, C3'I |
| 3'I | 4.43, m | H2'I, H4'I | 70.6/70.4 | C1'I, C4'I, C2'I, C5'I |
| 4'I | 4.39, m | H3'I, H5'I | 85.3 | C3'I, C5'I |
| 5'I | 4.21/4.18, m | H4'I | 65.2 | C4'I, C3'I |

## References

1. Murray, C. J. L.; Ikuta, K. S.; Sharara, F.; Swetschinski, L.; Robles Aguilar, G.; Gray, A.; Han, C.; Bisignano, C.; Rao, P.; Wool, E., et al. Global burden of bacterial antimicrobial resistance in 2019: a systematic analysis. The Lancet 2022, 399 (10325), 629–655.

2. Hatfull, G. F.; Dedrick, R. M.; Schooley, R. T. Phage Therapy for Antibiotic-Resistant Bacterial Infections. Annual Review of Medicine 2022, 73 (Volume 73, 2022), 197–211.

3. Oechslin, F. Resistance Development to Bacteriophages Occurring during Bacteriophage Therapy. Viruses 2018, 10 (7), 351.

4. Georjon, H.; Bernheim, A. The highly diverse antiphage defence systems of bacteria. Nature Reviews Microbiology 2023, 21 (10), 686–700.

5. Oromí-Bosch, A.; Antani, J. D.; Turner, P. E. Developing Phage Therapy That Overcomes the Evolution of Bacterial Resistance. Annual Review of Virology 2023, 10 (Volume 10, 2023), 503–524.

6. Bleriot, I.; Pacios, O.; Blasco, L.; Fernández-García, L.; López, M.; Ortiz-Cartagena, C.; Barrio-Pujante, A.; García-Contreras, R.; Pirnay, J.-P.; Wood, T. K., et al. Improving phage therapy by evasion of phage resistance mechanisms. JAC-Antimicrobial Resistance 2024, 6 (1).

7. Doron, S.; Melamed, S.; Ofir, G.; Leavitt, A.; Lopatina, A.; Keren, M.; Amitai, G.; Sorek, R. Systematic discovery of antiphage defense systems in the microbial pangenome. Science 2018, 359 (6379), eaar4120.

8. Ofir, G.; Herbst, E.; Baroz, M.; Cohen, D.; Millman, A.; Doron, S.; Tal, N.; Malheiro, D. B. A.; Malitsky, S.; Amitai, G., et al. Antiviral activity of bacterial TIR domains via immune signalling molecules. Nature 2021, 600 (7887), 116–120.

9. Roberts, C. G.; Fishman, C. B.; Banh, D. V.; Marraffini, L. A. A bacterial TIR-based immune system senses viral capsids to initiate defense. bioRxiv 2024, 2024.07.29.605636.

10. van den Berg, D. F.; Costa, A. R.; Esser, J. Q.; Stanciu, I.; Geissler, J. Q.; Zoumaro-Djayoon, A. D.; Haas, P.-J.; Brouns, S. J. J. Bacterial homologs of innate eukaryotic antiviral defenses with anti-phage activity highlight shared evolutionary roots of viral defenses. Cell Host & Microbe 2024, 32 (8), 1427–1443.e8.

11. Rousset, F.; Osterman, I.; Scherf, T.; Falkovich, A. H.; Leavitt, A.; Amitai, G.; Shir, S.; Malitsky, S.; Itkin, M.; Savidor, A., et al. TIR signaling activates caspase-like immunity in bacteria. Science 2025, 387 (6733), 510–516.

12. Sabonis, D.; Avraham, C.; Lu, A.; Herbst, E.; Silanskas, A.; Leavitt, A.; Yirmiya, E.; Zaremba, M.; Amitai, G.; Kranzusch, P. J., et al. TIR domains produce histidine-ADPR conjugates as immune signaling molecules in bacteria. bioRxiv 2024, 2024.01.03.573942.

13. Leavitt, A.; Yirmiya, E.; Amitai, G.; Lu, A.; Garb, J.; Herbst, E.; Morehouse, B. R.; Hobbs, S. J.; Antine, S. P.; Sun, Z.-Y. J., et al. Viruses inhibit TIR gcADPR signalling to overcome bacterial defence. Nature 2022, 611 (7935), 326–331.

14. Manik, M. K.; Shi, Y.; Li, S.; Zaydman, M. A.; Damaraju, N.; Eastman, S.; Smith, T. G.; Gu, W.; Masic, V.; Mosaiab, T., et al. Cyclic ADP ribose isomers: Production, chemical structures, and immune signaling. Science 2022, 377 (6614), eadc8969.

15. Tamulaitiene, G.; Sabonis, D.; Sasnauskas, G.; Ruksenaite, A.; Silanskas, A.; Avraham, C.; Ofir, G.; Sorek, R.; Zaremba, M.; Siksnys, V. Activation of Thoeris antiviral system via SIR2 effector filament assembly. Nature 2024, 627 (8003), 431–436.

16. Shi, Y.; Masic, V.; Mosaiab, T.; Rajaratman, P.; Hartley-Tassell, L.; Sorbello, M.; Goulart, C. C.; Vasquez, E.; Mishra, B. P.; Holt, S., et al. Structural characterization of macro domain– containing Thoeris antiphage defense systems. Science Advances 2024, 10 (26), eadn3310.

17. Storms, Z. J.; Teel, M. R.; Mercurio, K.; Sauvageau, D. The Virulence Index: A Metric for Quantitative Analysis of Phage Virulence. PHAGE 2020, 1 (1), 27–36.

18. Essuman, K.; Summers, D. W.; Sasaki, Y.; Mao, X.; Yim, A. K. Y.; DiAntonio, A.; Milbrandt, J. TIR Domain Proteins Are an Ancient Family of NAD^+^-Consuming Enzymes. Current Biology 2018, 28 (3), 421–430.e4.

19. Shi, Y.; Kerry, P. S.; Nanson, J. D.; Bosanac, T.; Sasaki, Y.; Krauss, R.; Saikot, F. K.; Adams, S. E.; Mosaiab, T.; Masic, V., et al. Structural basis of SARM1 activation, substrate recognition, and inhibition by small molecules. Molecular Cell 2022, 82 (9), 1643–1659.e10.

20. Bratkowski, M.; Burdett, T. C.; Danao, J.; Wang, X.; Mathur, P.; Gu, W.; Beckstead, J. A.; Talreja, S.; Yang, Y.-S.; Danko, G., et al. Uncompetitive, adduct-forming SARM1 inhibitors are neuroprotective in preclinical models of nerve injury and disease. Neuron 2022, 110 (22), 3711–3726.e16.

21. Neuts, A.-S.; Berkhout, H. J.; Hartog, A.; Goosen, J. H. M. Bacteriophage therapy cures a recurrent Enterococcus faecalis infected total hip arthroplasty? A case report. Acta Orthopaedica 2021, 92 (6), 678–680.

22. Köhler, T.; Luscher, A.; Falconnet, L.; Resch, G.; McBride, R.; Mai, Q.-A.; Simonin, J. L.; Chanson, M.; Maco, B.; Galiotto, R., et al. Personalized aerosolised bacteriophage treatment of a chronic lung infection due to multidrug-resistant Pseudomonas aeruginosa. Nature Communications 2023, 14 (1), 3629.

23. Engeman, E.; Freyberger, H. R.; Corey, B. W.; Ward, A. M.; He, Y.; Nikolich, M. P.; Filippov, A. A.; Tyner, S. D.; Jacobs, A. C. Synergistic Killing and Re-Sensitization of Pseudomonas aeruginosa to Antibiotics by Phage-Antibiotic Combination Treatment. Pharmaceuticals 2021, 14 (3), 184.

24. Tomich, P. K.; An, F. Y.; Damle, S. P.; Clewell, D. B. Plasmid-Related Transmissibility and Multiple Drug Resistance in Streptococcus faecalis subsp. zymogenes Strain DS16. Antimicrobial Agents and Chemotherapy 1979, 15 (6), 828–830.

25. Athukoralage, J. S.; White, M. F. Cyclic Nucleotide Signaling in Phage Defense and Counter-Defense. Annual Review of Virology 2022, 9 (Volume 9, 2022), 451–468.

26. Kazlauskiene, M.; Kostiuk, G.; Venclovas, Č.; Tamulaitis, G.; Siksnys, V. A cyclic oligonucleotide signaling pathway in type III CRISPR-Cas systems. Science 2017, 357 (6351), 605–609.

27. Niewoehner, O.; Garcia-Doval, C.; Rostøl, J. T.; Berk, C.; Schwede, F.; Bigler, L.; Hall, J.; Marraffini, L. A.; Jinek, M. Type III CRISPR–Cas systems produce cyclic oligoadenylate second messengers. Nature 2017, 548 (7669), 543–548.

28. Stella, G.; Marraffini, L. Type III CRISPR-Cas: beyond the Cas10 effector complex. Trends in Biochemical Sciences 2024, 49 (1), 28–37.

29. Cohen, D.; Melamed, S.; Millman, A.; Shulman, G.; Oppenheimer-Shaanan, Y.; Kacen, A.; Doron, S.; Amitai, G.; Sorek, R. Cyclic GMP–AMP signalling protects bacteria against viral infection. Nature 2019, 574 (7780), 691–695.

30. Millman, A.; Melamed, S.; Amitai, G.; Sorek, R. Diversity and classification of cyclic-oligonucleotide-based anti-phage signalling systems. Nature Microbiology 2020, 5 (12), 1608–1615.

31. Tal, N.; Morehouse, B. R.; Millman, A.; Stokar-Avihail, A.; Avraham, C.; Fedorenko, T.; Yirmiya, E.; Herbst, E.; Brandis, A.; Mehlman, T., et al. Cyclic CMP and cyclic UMP mediate bacterial immunity against phages. Cell 2021, 184 (23), 5728–5739.e16.

32. Loenen, W. A. M.; Dryden, D. T. F.; Raleigh, E. A.; Wilson, G. G.; Murray, N. E. Highlights of the DNA cutters: a short history of the restriction enzymes. Nucleic Acids Research 2013, 42 (1), 3–19.

33. Hille, F.; Richter, H.; Wong, S. P.; Bratovič, M.; Ressel, S.; Charpentier, E. The Biology of CRISPR-Cas: Backward and Forward. Cell 2018, 172 (6), 1239–1259.

34. Li, J.; Cheng, R.; Wang, Z.; Yuan, W.; Xiao, J.; Zhao, X.; Du, X.; Xia, S.; Wang, L.; Zhu, B., et al. Structures and activation mechanism of the Gabija anti-phage system. Nature 2024, 629 (8011), 467–473.

35. Antine, S. P.; Johnson, A. G.; Mooney, S. E.; Leavitt, A.; Mayer, M. L.; Yirmiya, E.; Amitai, G.; Sorek, R.; Kranzusch, P. J. Structural basis of Gabija anti-phage defence and viral immune evasion. Nature 2024, 625 (7994), 360–365.

36. Tuck, O. T.; Adler, B. A.; Armbruster, E. G.; Lahiri, A.; Hu, J. J.; Zhou, J.; Pogliano, J.; Doudna, J. A. Hachiman is a genome integrity sensor. bioRxiv 2024, 2024.02.29.582594.

37. Hu, H.; Hughes, T. C. D.; Popp, P. F.; Roa-Eguiara, A.; Martin, F. J. O.; Rutbeek, N. R.; Hendriks, I. A.; Payne, L. J.; Yan, Y.; Sousa, V. K. d., et al. Structure and mechanism of Zorya anti-phage defense system. bioRxiv 2023, 2023.12.18.572097.

38. Arkin, Michelle R.; Tang, Y.; Wells, James A. Small-Molecule Inhibitors of Protein-Protein Interactions: Progressing toward the Reality. Chemistry & Biology 2014, 21 (9), 1102–1114.

39. Radaeva, M.; Ton, A.-T.; Hsing, M.; Ban, F.; Cherkasov, A. Drugging the ‘undruggable’. Therapeutic targeting of protein–DNA interactions with the use of computer-aided drug discovery methods. Drug Discovery Today 2021, 26 (11), 2660–2679.

40. Maji, B.; Gangopadhyay, S. A.; Lee, M.; Shi, M.; Wu, P.; Heler, R.; Mok, B.; Lim, D.; Siriwardena, S. U.; Paul, B., et al. A High-Throughput Platform to Identify Small-Molecule Inhibitors of CRISPR-Cas9. Cell 2019, 177 (4), 1067–1079.e19.

41. Lim, D.; Zhou, Q.; Cox, K. J.; Law, B. K.; Lee, M.; Kokkonda, P.; Sreekanth, V.; Pergu, R.; Chaudhary, S. K.; Gangopadhyay, S. A., et al. A general approach to identify cell-permeable and synthetic anti-CRISPR small molecules. Nature Cell Biology 2022, 24 (12), 1766–1775.

42. Costa, A. R.; van den Berg, D. F.; Esser, J. Q.; Muralidharan, A.; van den Bossche, H.; Bonilla, B. E.; van der Steen, B. A.; Haagsma, A. C.; Fluit, A. C.; Nobrega, F. L., et al. Accumulation of defense systems in phage-resistant strains of Pseudomonas aeruginosa. Science Advances 2024, 10 (8), eadj0341.

43. Houte, S. v.; Buckling, A.; Westra, E. R. Evolutionary Ecology of Prokaryotic Immune Mechanisms. Microbiology and Molecular Biology Reviews 2016, 80 (3), 745–763.

44. Puigbò, P.; Makarova, K. S.; Kristensen, D. M.; Wolf, Y. I.; Koonin, E. V. Reconstruction of the evolution of microbial defense systems. BMC Evolutionary Biology 2017, 17 (1), 94.

45. Bernheim, A.; Sorek, R. The pan-immune system of bacteria: antiviral defence as a community resource. Nature Reviews Microbiology 2020, 18 (2), 113–119.

46. Zang, Z.; Zhang, C.; Park, K. J.; Schwartz, D. A.; Podicheti, R.; Lennon, J. T.; Gerdt, J. P. Bacterium secretes chemical inhibitor that sensitizes competitor to bacteriophage infection. *bioRxiv* 2024, 2024.01.31.578241.

47. Zang, Z.; Park, K. J.; Gerdt, J. P. A metabolite produced by gut microbes represses phage infections in *Vibrio cholerae*. ACS Chemical Biology 2022, 17 (9), 2396–2403.

48. Wang, D.-G.; Niu, L.; Lin, Z.-M.; Wang, J.-J.; Gao, D.-F.; Sui, H.-Y.; Li, Y.-Z.; Wu, C. The Discovery and Biosynthesis of Nicotinic Myxochelins from an Archangium sp. SDU34. Journal of Natural Products 2021, 84 (10), 2744–2748.

49. Frank, N. A.; Széles, M.; Akone, S. H.; Rasheed, S.; Hüttel, S.; Frewert, S.; Hamed, M. M.; Herrmann, J.; Schuler, S. M. M.; Hirsch, A. K. H., et al. Expanding the Myxochelin Natural Product Family by Nicotinic Acid Containing Congeners. Molecules 2021, 26 (16), 4929.

50. Wu, Y.; Garushyants, S. K.; van den Hurk, A.; Aparicio-Maldonado, C.; Kushwaha, S. K.; King, C. M.; Ou, Y.; Todeschini, T. C.; Clokie, M. R. J.; Millard, A. D., et al. Bacterial defense systems exhibit synergistic anti-phage activity. Cell Host & Microbe 2024, 32 (4), 557–572.e6.

51. Yasbin, R. E.; Wilson, G. A.; Young, F. E. Transformation and transfection in lysogenic strains of Bacillus subtilis: evidence for selective induction of prophage in competent cells. Journal of Bacteriology 1975, 121 (1), 296–304.

52. Guérout-Fleury, A.-M.; Frandsen, N.; Stragier, P. Plasmids for ectopic integration in Bacillus subtilis. Gene 1996, 180 (1), 57–61.

53. Huo, W.; Adams, H. M.; Zhang, M. Q.; Palmer, K. L. Genome Modification in Enterococcus faecalis OG1RF Assessed by Bisulfite Sequencing and Single-Molecule Real-Time Sequencing. Journal of Bacteriology 2015, 197 (11), 1939–1951.

54. Thurlow, L. R.; Thomas, V. C.; Hancock, L. E. Capsular Polysaccharide Production in Enterococcus faecalis and Contribution of CpsF to Capsule Serospecificity. Journal of Bacteriology 2009, 191 (20), 6203–6210.

55. Choi, K.-H.; Schweizer, H. P. mini-Tn7 insertion in bacteria with single attTn7 sites: example Pseudomonas aeruginosa. Nature Protocols 2006, 1 (1), 153–161.

56. Gibson, D. G.; Young, L.; Chuang, R.-Y.; Venter, J. C.; Hutchison, C. A.; Smith, H. O. Enzymatic assembly of DNA molecules up to several hundred kilobases. Nature Methods 2009, 6 (5), 343–345.

57. Piotto, M.; Saudek, V.; Sklenář, V. Gradient-tailored excitation for single-quantum NMR spectroscopy of aqueous solutions. Journal of Biomolecular NMR 1992, 2 (6), 661–665.

58. Sklenar, V.; Piotto, M.; Leppik, R.; Saudek, V. Gradient-Tailored Water Suppression for 1H-15N HSQC Experiments Optimized to Retain Full Sensitivity. Journal of Magnetic Resonance, Series A 1993, 102 (2), 241–245.

59. Mayer, M.; Meyer, B. Characterization of Ligand Binding by Saturation Transfer Difference NMR Spectroscopy. Angewandte Chemie International Edition 1999, 38 (12), 1784–1788.

60. Loyo, C. L.; Grossman, A. D. A phage-encoded counter-defense inhibits an NAD-degrading anti-phage defense system. *bioRxiv* 2024, 2024.12.23.630042.

61. Le, S.; Wei, L.; Wang, J.; Tian, F.; Yang, Q.; Zhao, J.; Zhong, Z.; Liu, J.; He, X.; Zhong, Q., et al. Bacteriophage protein Dap1 regulates evasion of antiphage immunity and Pseudomonas aeruginosa virulence impacting phage therapy in mice. Nature Microbiology 2024, 9 (7), 1828–1841.

## References for supplementary tables and figures

1. Ho, K.; Huo, W.; Pas, S.; Dao, R.; Palmer, K. L. Loss-of-Function Mutations in epaR Confer Resistance to &#x3d5;NPV1 Infection in Enterococcus faecalis OG1RF. Antimicrobial Agents and Chemotherapy 2018, 62 (10), 10.1128/aac.00758-18.

2. Chatterjee, A.; Johnson, C. N.; Luong, P.; Hullahalli, K.; McBride, S. W.; Schubert, A. M.; Palmer, K. L.; Carlson, P. E.; Duerkop, B. A. Bacteriophage Resistance Alters Antibiotic-Mediated Intestinal Expansion of Enterococci. Infection and Immunity 2019, 87 (6), 10.1128/iai.00085-19.

3. Huo, W.; Adams, H. M.; Zhang, M. Q.; Palmer, K. L. Genome Modification in Enterococcus faecalis OG1RF Assessed by Bisulfite Sequencing and Single-Molecule Real-Time Sequencing. Journal of Bacteriology 2015, 197 (11), 1939–1951.

